# Chromatin stability safeguards mitochondrial homeostasis and prevents mTORC1 hyperactivation

**DOI:** 10.64898/2025.12.09.693285

**Authors:** Vinoth Sigamani, Mohd Yousuf, Daniel L. Johnson, Brian D. Strahl, R. Nicholas Laribee

**Affiliations:** Department of Pathology and Laboratory Medicine, College of Medicine and the Center for Cancer Research, University of Tennessee Health Science Center, Memphis, TN, United States of America; Molecular Bioinformatics Core and the University of Tennessee Health Science Center Office of Research, University of Tennessee Health Science Center, Memphis, TN, United States of America; Department of Biochemistry and Biophysics, and the Lineberger Comprehensive Cancer Center, University of North Carolina at Chapel Hill, Chapel Hill, NC, United States of America

**Keywords:** Histone stability, metabolic stress, Set2, mTORC1, chromatin, ROS, mitochondrial homeostasis

## Abstract

Cells dynamically regulate chromatin in response to nutrient flux which promotes the transcriptional changes necessary for adaptation. The mechanistic target of rapamycin complex 1 (mTORC1) kinase integrates nutrient signaling with chromatin regulation, yet whether chromatin stability feeds back to mTORC1 activation and stress adaption remains unknown. We previously identified histone H3 at lysine 37 (H3K37) as essential for the response to mTORC1 stress such that mutation of H3K37 to alanine (H3K37A) causes cell death upon mTORC1 inhibition. Herein, we show that H3K37-dependent chromatin stability prevents proteasome-mediated histone degradation, restricts mTORC1 signaling, and safeguards mitochondrial homeostasis during mTORC1 stress. Genetic interaction analyses reveal that H3K37A combined with mutants that destabilize chromatin, including loss of the Set2 H3K36 methyltransferase, Rpd3S histone deacetylase, or multiple histone deposition pathways, causes synthetic lethality when mTORC1 is inhibited. Transcriptome analysis indicates that H3K37A misregulates the mitochondrial transcriptome during mTORC1 stress, which increases mitochondrial reactive oxygen species (ROS) and triggers lethal mitochondrial retrograde signaling. Inactivation of retrograde signaling, or ROS neutralization, rescues viability of H3K37A and chromatin stability mutants during mTORC1 stress. These findings establish chromatin stability as a key safeguard that restrains mTORC1 activity and prevents toxic mitochondrial stress during metabolic adaptation.

## Introduction

Eukaryotic cells dynamically adjust their growth and metabolism in response to changes in nutrient availability, energy status, and environmental stress^1^. A central regulator of this adaptive response is the mechanistic target of rapamycin complex 1 (mTORC1), a conserved kinase complex that integrates nutrient inputs with downstream control of transcriptional regulation, protein synthesis, metabolism, and autophagy inhibition required for cell growth and proliferation^1,2^. In all eukaryotes, coordination of mTORC1 activity with the availability of nutrient resources is essential to prevent deregulated growth or activation of cell stress responses that compromise organismal fitness^3–5^.

In budding yeast, the mTORC1 complex is a serine/threonine kinase comprised of either the Tor1 or Tor2 kinase subunits along with the additional Lst8, Kog1, and Tco89 subunits^6,7^. In yeast, the vacuole (homologous to the mammalian lysosome) is a degradative compartment that stores amino acids and other metabolites^8^. mTORC1 is recruited to the vacuole surface via Kog1 and Tco89-mediated interactions with the Rag GTPase containing EGO complex that consists of the Rag GTPases Gtr1 and Gtr2, and the additional Ego1-3 subunits^2,9,10^. Vacuole amino acids activate mTORC1 by promoting Gtr1 loading with GTP and Gtr2 loading with GDP. Interaction of mTORC1 with EGO and the additional vacuole V-ATPase complex, which is the resident vacuole H^+^-pump that acidifies the vacuole, results in full mTORC1 activation^11^. Once activated, mTORC1 phosphorylates a complex network of direct substrates that promote anabolism and cell proliferation while inhibiting nutrient stress responses, including autophagy^2,12–14^. Among the multiple downstream kinases phosphorylated by mTORC1 is the Ypk3 kinase that in turn phosphorylates ribosomal protein S6 (S6ph) ^15^. Emerging evidence indicates that mTORC1 also directly controls gene transcription in the nucleus, and it is a known regulator of epigenetic signaling ^16,17^. However, the underlying mechanisms by which mTORC1 controls transcription and the epigenome remain poorly defined. Deciphering this regulation will ultimately be essential for determining how mTORC1 dysregulation causes disease.

The regulation of chromatin structure and histone post-translational modifications (PTMs), collectively referred to as epigenetic regulation, controls gene expression and genome stability, and this epigenetic regulation is highly intertwined with cellular metabolism^18,19^. The abundance of many histone PTMs that affect gene expression, including histone methylation and acetylation, is controlled by the availability of their metabolic donor molecules (S-adenosylmethionine (SAM) for histone methylation and acetyl-CoA for histone acetylation), which orchestrate chromatin accessibility and transcriptional output with the cellular metabolic state^20^. Importantly, recent studies suggest that chromatin not only responds to metabolic cues, but that it also can signal in reverse to potentially modulate nutrient-responsive signaling pathways such as mTORC1^21,22^. These studies suggest that reciprocal communication may exist between chromatin and mTORC1, yet the mechanisms explaining such bidirectional signaling have yet to be defined.

We previously performed a chemical genetic screen utilizing the mTORC1-specific inhibitor rapamycin against a library of histone H3 and H4 (H3/H4) mutants to identify functional relationships between mTORC1 and the epigenome^23^. This screen found that mTORC1 exhibits strong genetic interactions with multiple histone H3 N-terminal residues, many of which are sites of PTM. Notably, we found that mutation of H3K37, which is located at the nucleosomal DNA entry and exit point^24^, to alanine (H3K37A) causes cytotoxicity upon mTORC1 inhibition^21,23^. H3K37 is adjacent to H3K36, which is methylated by the Set2 methyltransferase required for maintaining transcriptional fidelity and chromatin stability during gene transcription^25,26^. Importantly, both the H3K37A mutant, as well as cells lacking Set2 (*set2Δ*), exhibit elevated mTORC1 signaling^21,22^, suggesting that the H3K36/K37 di-lysine motif may contribute to chromatin-mediated mTORC1 regulation. Set2 catalyzes mono-, di-, and tri-methylation of H3K36 (H3K36me1/2/3) via its conserved SET domain, whereas its conserved C-terminal Set2-Rpb1 interacting (SRI) domain binds to the dually phosphorylated serine 2 and serine 5 residues of the RNA polymerase II (Pol II) C-terminal domain (CTD)^25–27^. Set2 binding to Pol II promotes targeting of H3K36me1/2/3 to transcribed chromatin, which maintains transcriptional fidelity and chromatin integrity through the recruitment of epigenetic reader proteins that selectively bind the H3K36me2/3 modifications^26^. Specific readers of H3K36me2/3 include the Eaf3 subunit of the Rpd3S histone deacetylase (HDAC) complex^28–30^, the Ioc4 subunit of the Isw1b chromatin remodeling complex^31^, and the Pdp3 subunit of the NuA3 histone acetyltransferase complex^32^. To date, no known reader protein has been identified that selectively binds H3K36me1. Importantly, H3K36me2/3 also maintains chromatin stability during transcription by suppressing histone replacement mediated by histone chaperone proteins such as Asf1^33^.

In this study, we used the H3K37A mutation as a tool to define how chromatin stability constrains mTORC1 signaling and supports cellular adaptation to mTORC1 stress. We show that H3K37A destabilizes histones, leading to proteasome-dependent histone degradation that is associated with mTORC1 hyperactivation. Genetic and biochemical analyses reveal that Set2-mediated H3K36me2/3 and the Rpd3S HDAC complex cooperates with H3K37 to prevent chromatin instability and sustain viability during mTORC1 stress. Transcriptome analyses indicates that mTORC1 inhibition in H3K37A selectively deregulates genes controlling mitochondrial function and metabolism, driving mitochondrial reactive oxygen species (ROS) and cytotoxic mitochondrial retrograde signaling. Inhibition of mitochondrial retrograde signaling or ROS neutralization fully rescues H3K37A viability under mTORC1 stress. Compromising additional chromatin assembly pathways in the H3K37A background triggers a similar lethal mitochondrial retrograde program, underscoring that maintaining chromatin stability prevents toxic activation of this pathway during mTORC1 stress. We also find that combining H3K37A with either Set2 loss or chromatin assembly defects further hyperactivates mTORC1 activity, supporting a causal link between histone degradation and mTORC1 deregulation. These findings establish chromatin stability as a key safeguard that restrains mTORC1 signaling, preserves mitochondrial homeostasis, and shapes metabolic stress adaptation.

## Results

### H3K37 prevents histone destabilization and sensitivity to mTORC1 stress

We previously found that mutation of H3K37 to alanine (H3K37A) resulted in a loss of HMGB protein binding to chromatin, and it also increased both mTORC1 signaling and sensitivity to mTORC1 stress ^21,23^. H3K37 is located at the nucleosome DNA entry/exit point, and a recent nucleosome crystal structure suggests that H3K37 may contact nucleosomal DNA to help stabilize epigenetic reader binding to the adjacent methylated H3K36 residue (**Fig. 1A**)^34^. To delineate the functional relationship between H3K37 and the mTORC1 pathway, we initially tested the idea that the H3K37A hypersensitivity to mTORC1 stress might be due to a histone defect. H3 wild-type (H3WT) and H3K37A were transformed with either a control vector or a vector expressing all four core histone genes. Cells were then spotted to control media or media containing the mTORC1 inhibitor rapamycin. While H3K37A vector control cells exhibited a profound growth defect compared to H3WT, histone expression rescued H3K37A growth under mTORC1 stress **(Supplementary Fig. S1A).** Intriguingly, immunoblot (IB) analysis of both soluble and denatured cell extracts revealed that histone protein levels were reduced in the H3K37A vector control relative to the H3WT vector control, and this histone deficiency was rescued by the histone expression vector (**Fig. 1B-E and Supplementary Fig. S1B-E)**. These data indicate that H3K37A causes both a global reduction in histone levels, and an increased sensitivity to mTORC1 stress, that can be rescued by histone overexpression.

**Figure 1.**
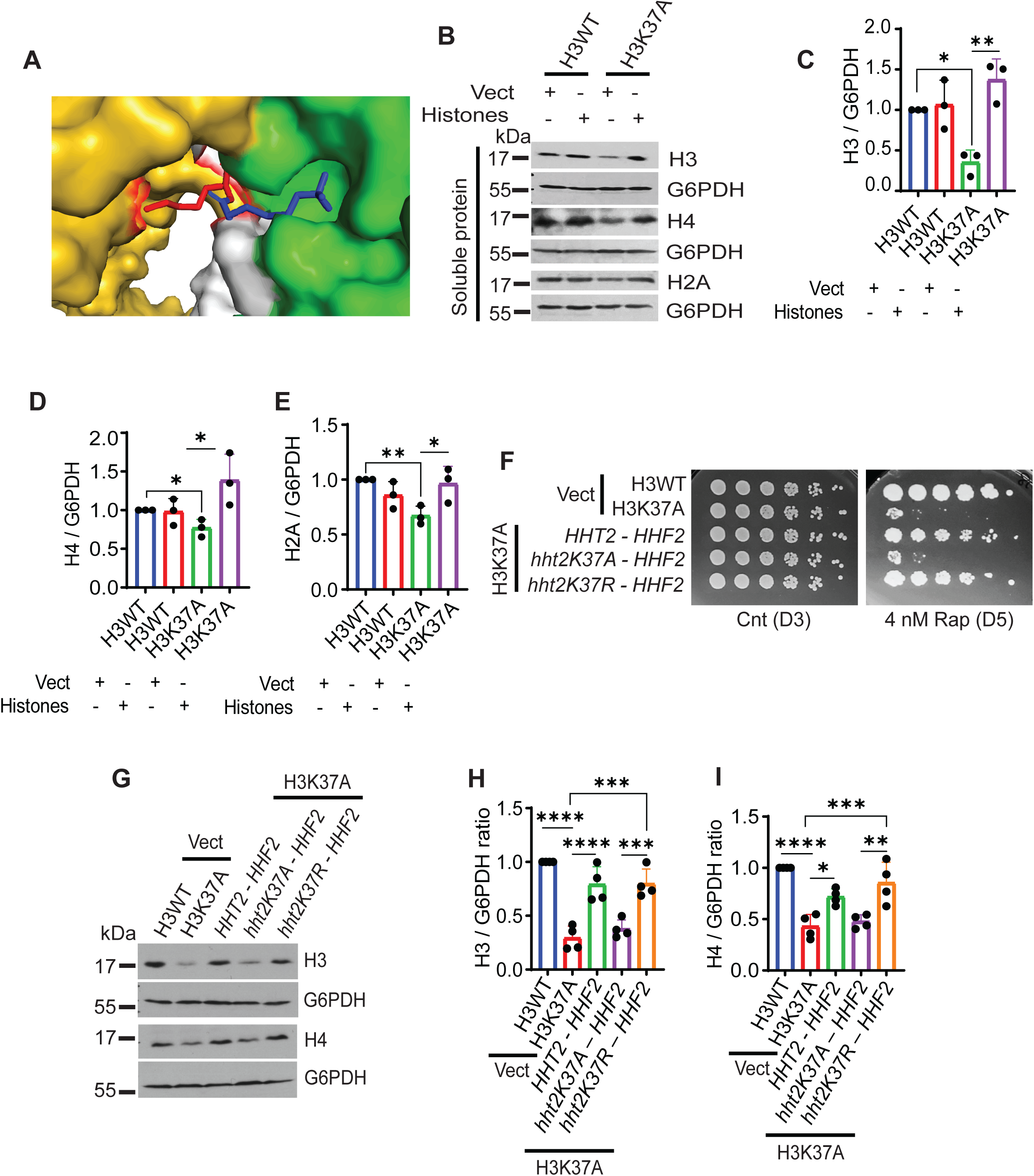
The H3K37A mutant reduces histone expression and causes lethality during mTORC1 stress. **(A)** The crystal structure of nucleosome-Rpd3S interaction. Red-H3K37; blue-H3K36me3; yellow– DNA; white-histones; green– Eaf3. Structure is PDB ID 8TOF. **(B)** H3WT and H3K37A were transformed with a control vector or a vector expressing all four core histone genes. Soluble protein extracts were prepared and analyzed by IB with the indicated antibodies. **(C-E)** Quantification of histones H3, H4, and H2A and normalized to G6PDH signal. n=3; *\*p<0.05*, *\*\*p<0.01*. **(F)** The H3K37A cells transformed with the *HHT2-HHF2, hht2K37A-HHF2*, or *hht2K37R-HHF2* plasmid, along with the control vector in H3WT and H3K37A, were diluted 5-fold and spotted onto the indicated plate for 3-5 days. **(G)** IBs for the total histone H3 and H4 for the cells in F. **(H-I)** The graph indicates IB band intensity of H3, H4, and was normalized to G6PDH. n=4; *\*p<0.05*, *\*\*p<0.01*, *\*\*\*p<0.001*, *\*\*\*\*p<0.0001*.

We next confirmed that the H3K37A mutation was solely responsible for the mTORC1 stress sensitivity and histone reduction. H3WT and H3K37A were transformed with a control vector, while H3K37A was also transformed with a vector expressing either the wild-type *HHT2-HHF2* histone H3-H4 gene locus, an H3K37A mutation (*hht2K37A-HHF2*) that disrupts both the size and charge of the native H3K37 residue, or an H3K37R change (*hht2K37R-HHF2*) that restores charge and size at this position. Cells were then spotted to control media or rapamycin media. None of the strains exhibited growth defects in control media, whereas the H3K37A vector control cells were severely growth-impaired under rapamycin-induced mTORC1 stress (**Fig. 1F**). The *HHT2-HHF2* and *hht2K37R-HHF2* expression vectors rescued this growth deficiency, while *hht2K37A-HHF2* did not (**Fig. 1F**). Additionally, IB analysis of H3K37A cells expressing either *HHT2-HHF2* or *hht2K37R-HHF2* revealed that histone protein levels were restored to those of H3WT, while histone levels in *hht2K37A-HHF2* expressing cells were similar to the H3K37A control (**Fig. 1G-I**). Our previous report that employed individual yeast strains expressing these histone H3 mutants integrated into the genome also found that H3WT, H3K37R, and H3K37Q restored growth during mTORC1 stress, whereas H3K37A did not ^21^. Collectively, these findings demonstrate that the single H3K37A mutation mediates hypersensitivity to mTORC1 stress and globally reduces histone levels.

Since *hht2K37A-HHF2* failed to restore histone levels, we next determined if H3K37A reduced histone gene expression to explain the lower histone protein levels. The H3WT and H3K37A strains express a single genomically integrated synthetic H3-H4 cassette^35^, while the endogenous *HHT1-HHF1* and *HHT2-HHF2* alleles have been deleted. Using *HHT1*-specific primers that recognize the synthetic H3 cassette, we analyzed *HHT1, HTA1,* and *HTB1* mRNA levels by quantitative real-time PCR (qRT-PCR) in H3WT and H3K37A expressing control vector, and also in H3K37A expressing either *HHT2-HHF2* or *hht2K37A-HHF2*. No differences in *HHT1, HTA1, and HTB1* mRNA levels were found between H3WT and either H3K37A control vector, *HHT2-HHF2,* or *hht2K37A-HHF2* expressing cells (**Supplementary Fig. S1F-H**), therefore indicating the histone H3 reduction by H3K37A is independent of mRNA changes. Additionally, no difference in *HHT2* or *hht2K37A* expression from the exogenous histone vector was detected in H3K37A (**Supplementary Fig. S1I**). Based on these data and additional results discussed below, we conclude that H3K37A reduces histone levels through an mRNA-independent mechanism.

Histones are synthesized during the S-phase where they are then incorporated into nascent chromatin^36^. We assessed whether cell cycle differences exist between H3WT and H3K37A that could explain the histone reduction in H3K37A. Asynchronous H3WT and H3K37A cultures were stained with SYTOX Green and analyzed by flow cytometry, which revealed that H3WT and H3K37A exhibited nearly identical cell cycle profiles (**Fig. 2A and Supplementary Fig. S2A-B)**. Therefore, H3K37A inhibits histone expression independently of changes to its cell cycle. We next determined whether H3K37A enhanced histone degradation to account for the lower histone levels. Protein degradation occurs predominantly through either autophagy or the proteasome. We blocked autophagy in H3WT and H3K37A using the autophagy inhibitor phenylmethylsulfonyl fluoride (PMSF)^37^, and histone IB analysis revealed that autophagy inhibition failed to restore histone expression (**Fig. 2B-C**). However, proteasome inhibition in H3WT and H3K37A using the proteasome inhibitor MG132 restored H3K37A histone levels to those of H3WT (**Fig. 2D-F**)^38^. These data suggest that H3K37A increases histone instability and subsequent degradation by the proteasome. Since a reduction in histone stability may promote chromatin decompaction^39^, we analyzed global chromatin accessibility in H3WT and H3K37A by performing micrococcal nuclease (MNase) digestion. Even though H3K37A quantifiably reduces histone levels (**Fig. 1B-E, 1G, and Supplementary Fig. S1B-E**), no difference in chromatin accessibility was detectable between H3WT and H3K37A **(Supplementary Fig. S2C)**. In total, these findings indicate that while H3K37A increases histone instability and degradation by the proteasome, the histone reduction observed is insufficient to substantively alter global chromatin compaction.

**Figure 2.**
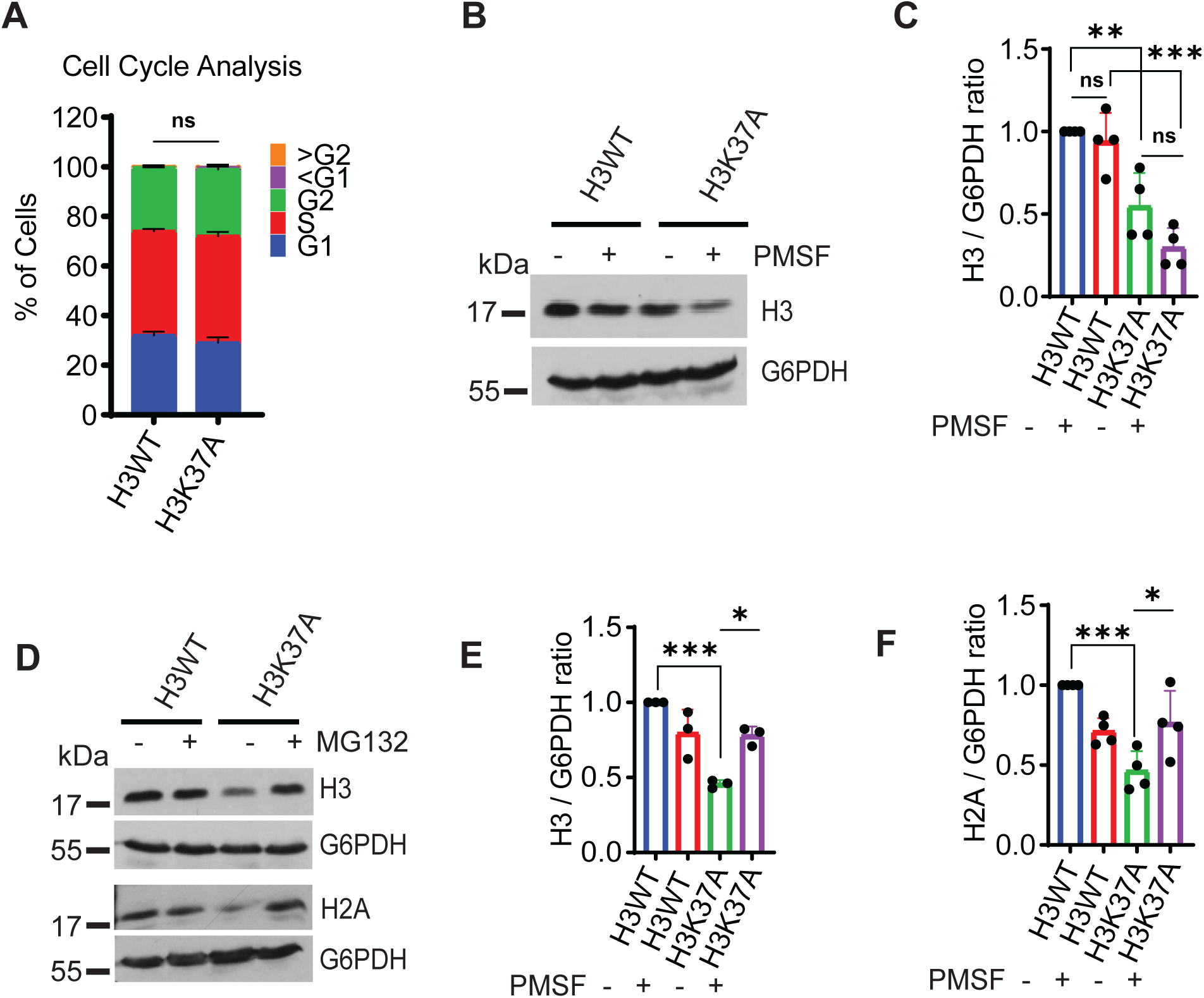
The H3K37A mutant causes histone instability and degradation via the proteasome. **(A)** Graphical representation of the cell percentage for each phase of the cell cycle in H3WT and H3K37A. n=3; ns-no significance. **(B)** The H3WT and H3K37A were treated with the autophagy inhibitor PMSF (1 mM) for four hours before analyzing histone H3 by IB. **(C)** Quantification of histone H3 levels normalized to G6PDH from **B**. n=4; *\*\*p<0.01*, *\*\*\*p<0.001*. **(D)** H3WT and H3K37A are treated with the proteosome inhibitor MG132 (20 µM) for two hours, and then histones H3 and H2A were analyzed by IB. **(E-F)** Quantification of histones H3 (**E**) or H2A (**F**) normalized to G6PDH. n=3-4; *\*p<0.05*, *\*\*\*p<0.001*.

### H3K37 and the Set2 methyltransferase act synergistically to sustain viability during mTORC1 stress

H3K37 is adjacent to H3K36 (**Fig. 1A**), which is methylated co-transcriptionally by the Set2 methyltransferase. We previously found that Set2 regulates mTORC1 signaling and the response to mTORC1 stress^22^. To assess if loss of Set2 in H3K37A increases sensitivity to mTORC1 stress, we combined a *set2Δ* into H3WT and H3K37A. Set2 is one of three histone H3 methyltransferases with critical roles in transcriptional control, so we also expanded our analysis to examine inactivating mutants of these additional histone H3 methyltransferases. H3WT and H3K37A strains were generated that either lack H3K4 methylation due to a Set1-containing COMPASS complex inactivating mutant (*cps50Δ*) or lack Dot1-dependent H3K79 methylation (*dot1Δ*)^40,41^. All strains were scored for their mTORC1 stress sensitivity by plating onto control media or media containing rapamycin. Neither the H3WT, nor the individual methyltransferase mutants in the H3WT background, affected mTORC1 stress sensitivity (**Fig. 3A**). As expected, the H3K37A mutant exhibited sensitivity to mTORC1 stress, and this effect was further exacerbated by *set2Δ* (**Fig. 3A**). Since the only known Set2 substrate is H3K36^26^, the increased mTORC1 stress sensitivity of H3K37A *set2Δ* indicates that Set2-mediated H3K36 methylation remains at least partially intact in H3K37A. Importantly, while the H3K37A *cps50Δ* caused only a negligible increase in mTORC1 stress sensitivity relative to H3K37A, the H3K37A *dot1Δ* modestly rescued the H3K37A growth defect (**Fig. 3A**). Although both COMPASS and Set2 were proposed to methylate H3K37^42^, the rescue of the H3K37A mTORC1 growth defect by either H3K37R (**Fig. 1F**) or H3K37Q^23^, combined with the negligible effect that H3K37A *cps50Δ* has (**Fig. 3A**), indicates that loss of H3K37 methylation is not responsible for the increased sensitivity to mTORC1 stress. Set2-mediated H3K36 methylation prevents nucleosome hyperacetylation and disruption of nucleosome stability during transcription^26^, while Dot1-mediated H3K79 methylation is associated with an open and transcriptionally active chromatin environment^43^. The opposing effects that Set2 and Dot1 loss have on the H3K37A mTORC1 stress response suggest that maintaining chromatin stability is essential for viability during mTORC1 stress.

**Figure 3.**
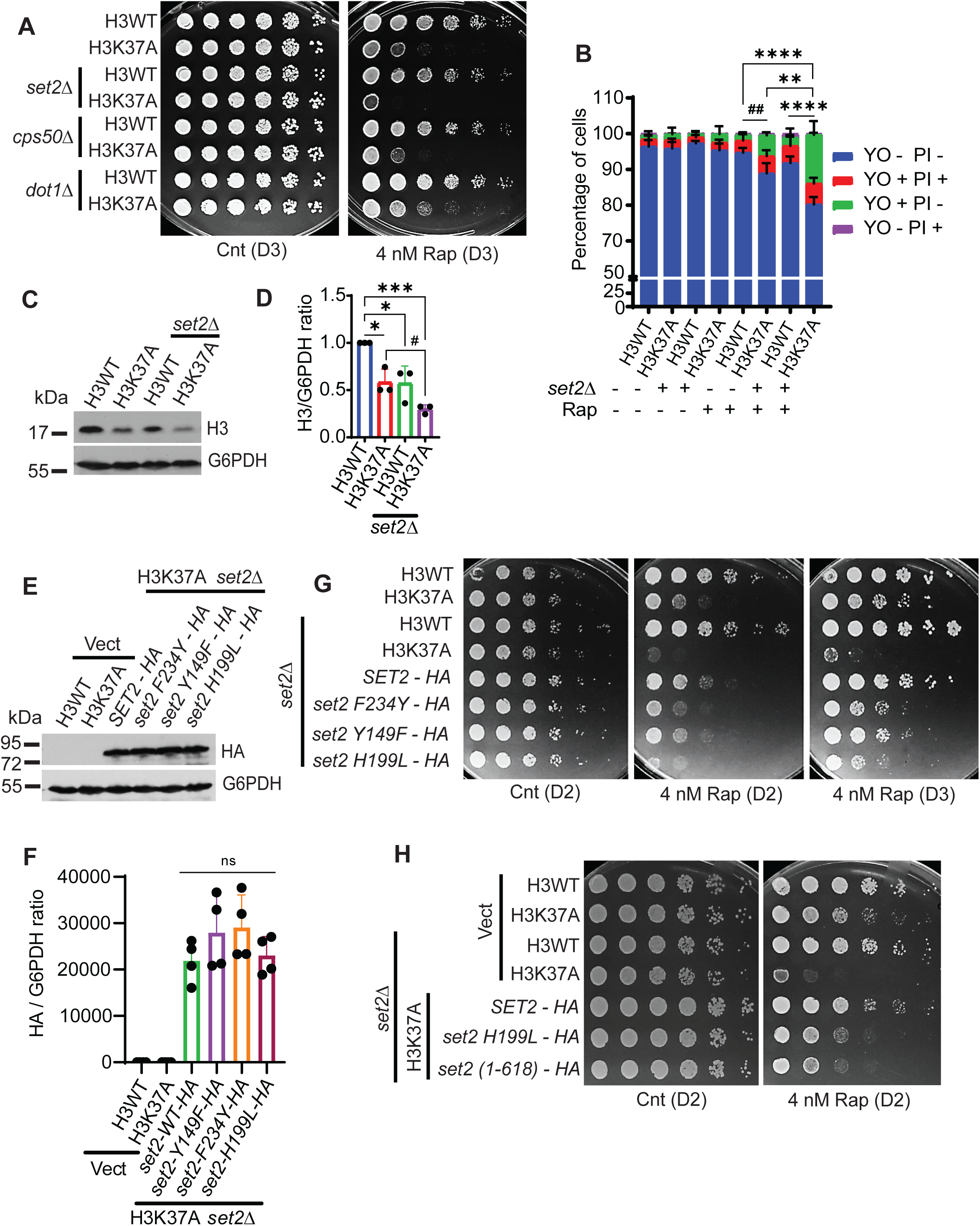
Set2 maintains viability in H3K37A during mTORC1 stress through methylation-dependent and -independent mechanisms. **(A)** The *set2Δ, cps50Δ,* and *dot1Δ* were diluted 5-fold and spotted onto the indicated plates for three days. **(B)** H3WT, H3K37A, and their *set2Δ* derivatives were mock-treated or 50 nM rapamycin-treated for six hours before YO-PRO-1 and PI staining and analysis by flow cytometry. The cell death percentage is represented in the graph. n=3; *\*\*p<0.01*, *\*\*\*\*p<0.0001* and *^##^p<0.01*. **(C)** IB analysis of histone H3 in H3WT, H3K37A, and their *set2Δ* derivatives. **(D)** Quantification of histone H3 normalized to G6PDH from **C**. n=3; *\*p<0.05*, *\*\*\*p<0.001* and *^#^p<0.05*. **(E)** Expression of wild-type Set2-HA and the indicated Set2 mutants in H3K37A *set2Δ*. **(F)** Quantification of Set2-HA and the indicated Set2-HA mutants normalized to G6PDH from **E**. n=4; ns-no significance. **(G)** Spot assay of the strains transformed with the control vector, or vectors expressing either wild-type Set2-HA or the indicated Set2 mutants. **(H)** As in **G,** except cells express either control vector, or vector expressing wild-type Set2, the catalytically inactive Set2 (*set2-H199L*), or the SRI domain-truncated Set2 mutant (*set2 (1-618)*). For **A**, **G**, and **H**, images were captured on the indicated day.

We previously found that the H3K37A growth defect during mTORC1 stress is due to cell death^21^. Since H3K37A *set2Δ* enhanced sensitivity to mTORC1 stress, we tested if it exhibited increased cell death upon mTORC1 inhibition. Asynchronous cultures of H3WT, H3K37A, and their *set2Δ* derivatives were mock-treated or treated with 50 nM rapamycin for six hours before staining with YO-PRO-1 (YO, stains early apoptotic cells) and propidium iodide (PI, stains late apoptotic and necrotic cells) and then analyzed by flow cytometry **(Fig. 3B and Supplementary Fig. S3A)**. No significant cell death was detected in the H3WT, H3K37A, and the *set2Δ* mutants under control conditions (**Fig. 3B**). Importantly, while rapamycin treatment did not affect either H3WT or H3WT *set2Δ*, it increased cell death in H3K37A as expected^21^, and this cytotoxic effect was enhanced further by H3K37A *set2Δ* (**Fig. 3B**). To determine if the increased cell death in H3K37A *set2Δ* was linked to a further reduction in histone levels, we quantified histone H3 levels in H3WT, H3K37A, and the *set2Δ* mutants. H3K37A reduced H3 expression as seen above (**Fig. 1**), while H3WT *set2Δ* also reduced H3 levels, and this histone reduction was decreased further by H3K37A *set2Δ* **(Fig. 3C-D**). These data demonstrate that loss of the Set2 pathway in H3K37A enhances mTORC1-stress induced cytotoxicity and further reduces histone protein levels. However, histone instability alone cannot be the sole cause of cell death under these conditions since the H3WT *set2Δ* has reduced histone expression but a normal response to mTORC1 stress.

Set2 is solely responsible for mono-, di-, and tri-methylation at H3K36 (H3K36me1/2/3)^26^. We next determined the role the specific H3K36 methylation states have in sustaining the viability of H3K37A *set2Δ* during mTORC1 stress. H3K37A *set2Δ* was transformed with control vector, or vectors expressing either HA-tagged *SET2* wild-type (*SET2-WT*), the *set2-Y149F* mutant that selectively maintains H3K36me3 without H3K36me1/2, the *set2-F234Y* that generates H3K36me1/2 but lacks H3K36me3, or the catalytic domain mutant *set2-H199L* that ablates almost all H3K36 methylation while retaining only minimal residual H3K36me1^44^. The Set2-WT and the different mutants were equivalently expressed in H3K37A **(Fig. 3E-F),** and they restored the expected H3K36 methylation states as previously reported **(Supplementary Fig. S3B)**^44^. Under mTORC1 stress, the *SET2-WT* rescued H3K37A *set2Δ* viability and growth back to that of H3K37A (**Fig. 3G**). Although less effective than Set2-WT, expressing either Set2-F234Y, Set2-Y149F, or Set2-H199L in H3K37A *set2Δ* rescued growth relative to H3K37A *set2Δ* vector control **(Fig. 3G).** Set2 is recruited to chromatin in part by binding to the phosphorylated C-terminal domain of Pol II through the Set2 C-terminal SRI domain. Set2 binding to the CTD then alleviates autoinhibition of Set2 catalytic activity to generate H3K36me2/3^45^. To test if Set2 requires Pol II binding for its role in the mTORC1 stress response, we transformed H3K37A *set2Δ* with either *SET2-WT-HA*, *set2(1-618)-HA* that lacks the SRI domain, the catalytically inactive *set2-H199L-HA*, or a control vector. Spot assays revealed that the *SET2-WT* rescued H3K37A *set2Δ* growth back to that of H3K37A control (**Fig. 3H**). Importantly, while less efficient than *SET2-WT-HA*, both *set2(1-618)-HA* and *set2-H199L-HA* partially rescued growth under mTORC1 stress **(Fig. 3H**). The SRI truncated Set2 does not bind Pol II or generate H3K36me3, likely due to autoinhibition of its SET domain^27,45^, while the *set2-H199L* generates only minimal residual H3K36me1 with no H3K36me2/3^44^. No epigenetic reader is known to selectively bind H3K36me1, and no functional role for H3K36me1 has ever been defined. Therefore, these results suggest that Set2 may provide both a methylation-dependent and methylation-independent role in maintaining H3K37A viability during mTORC1 stress that only partially depends upon Pol II binding.

We next addressed the functional consequence that loss of H3K36me2/3 demethylase activity (mediated by Rph1) in H3K37A has on the mTORC1 stress response^46^. A *rph1Δ* was introduced into H3WT and H3K37A, and then H3WT, H3K37A, and both the *set2Δ* and *rph1Δ* derivatives were scored on control and rapamycin media. Consistent with the above results, H3K37A *set2Δ* caused synthetic lethality upon mTORC1 stress, while the H3K37A *rph1Δ* resulted in a modest growth rescue (**Fig. 4A**). These results agree with the above data demonstrating that inhibition of H3K37A mTORC1-stress induced cytotoxicity partially depends on H3K36 methylation (**Fig. 3G**). H3K36me2/3 binds distinct epigenetic reader proteins to regulate chromatin structure during transcription, including the Eaf3 subunit of the Rpd3S histone deacetylase (HDAC) that binds H3K36me2/3^28–30^, the Ioc4 subunit of the Isw1b chromatin remodeler that binds H3K36me3^31^, and the Pdp3 subunit of the NuA3b HAT complex that also binds H3K36me3^32^. To identify the readers relevant to the H3K37A mTORC1 stress response, we individually deleted the genes encoding each reader subunit in H3WT and H3K37A and then scored each for growth on control or rapamycin plates. Loss of either Eaf3 or its associated HDAC subunit (Rpd3) phenocopied Set2 loss (**Fig. 4A**), indicating that Set2-dependent suppression of mTORC1 stress-induced cell death requires Rpd3S chromatin anchoring and histone deacetylation. Consistent with this concept, we found that H3K23ac levels (an Rpd3S substrate^47^) increased in H3K37A, H3WT *set2Δ*, and H3K37A *set2Δ* compared to H3WT **(Supplementary Fig. S3C-D)**. While H3WT *set2Δ* is expected to increase histone acetylation due to disrupted Rpd3S chromatin binding, these data suggest that H3K37A also may modestly reduce Rpd3S binding. This possibility is not unexpected since H3K37 interacts with DNA to potentially stabilize Eaf3-dependent nucleosome binding (**Fig. 1A**). Intriguingly, the H3K37A *set2Δ* mutant exhibits lower H3K23ac relative to H3WT *set2Δ* **(Supplementary Fig. S3C-D),** but the reason for this reduction is currently unclear. Analysis of the additional H3K36me2/3 readers revealed that H3K37A *ioc4Δ*, which disrupts anchoring of Isw1b to transcribed regions and impairs Isw1b-mediated nucleosome positioning and suppression of histone exchange^31^, resulted in a minor synthetic sick interaction compared to H3K37A (**Fig. 4B**). Importantly, we consistently observed that the H3K37A *pdp3Δ*, which impairs NuA3 histone acetyltransferase binding to transcribed genes^32^, rescued growth relative to H3K37A (**Fig. 4B**). Overall, these data suggest that H3K37A increases sensitivity to mTORC1 stress by impinging on pathways that sustain chromatin stability, including histone deacetylation.

**Figure 4.**
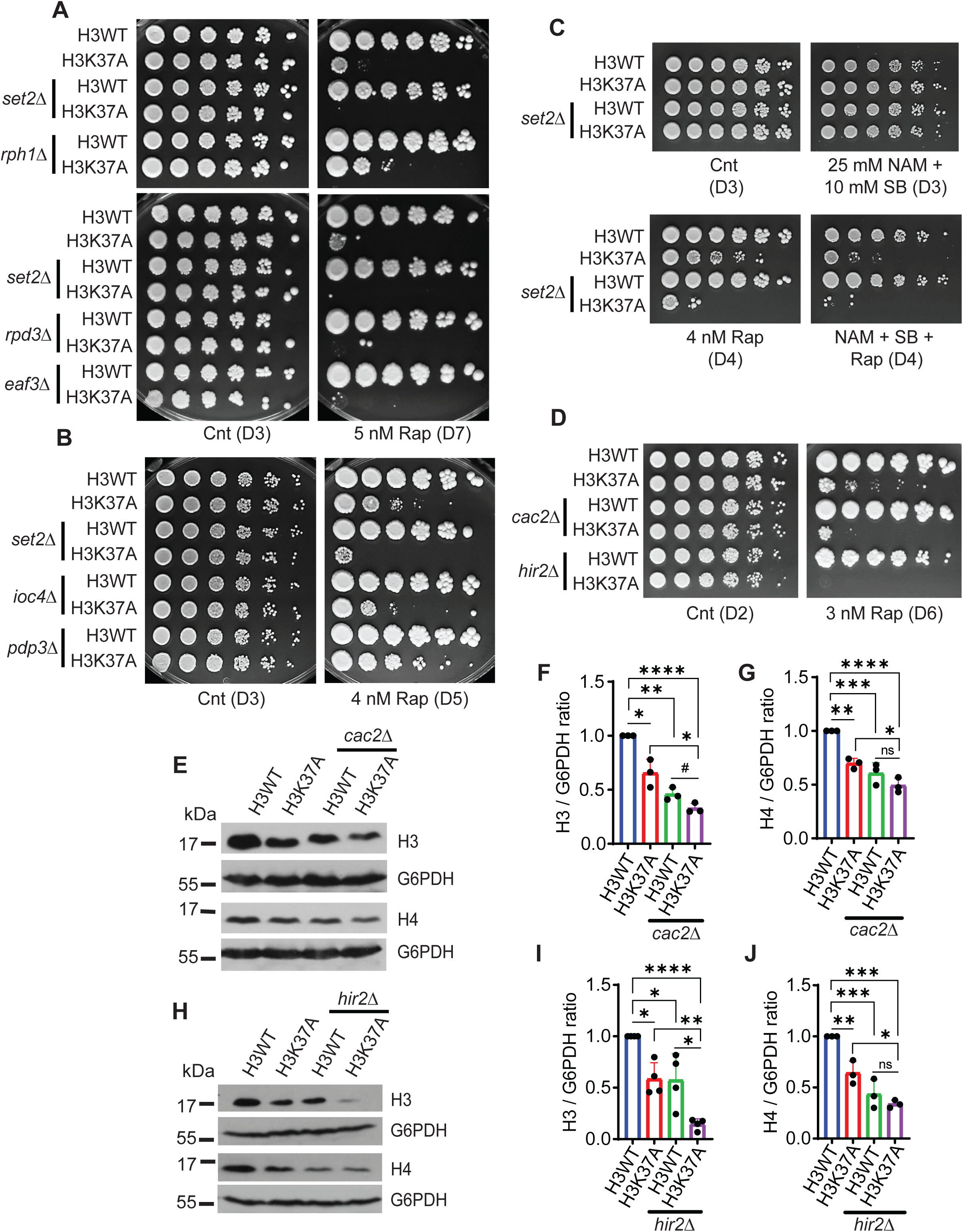
The Set2, HIR, and CAF-1 pathways prevent increased histone instability and mTORC1 stress in H3K37A. (A-B) The indicated strains were spotted onto control and rapamycin (4-5 nM) plates for 3-7 days. **(C)** H3WT, H3K37A, and their *set2Δ* derivatives cells were spotted onto control plates, or plates containing HDAC inhibitors (25 mM NAM and 10 mM SB), 4 nM rapamycin, or all inhibitors combined for 3-4 days. **(D)** The H3WT and H3K37A *cac2Δ* and *hir2Δ* mutants were serially diluted 5-fold and spotted onto the indicated plates for 2-6 days. **(E-G)** IB analysis and quantification of histones H3 and H4 in H3WT and H3K27A *cac2Δ*. Histone H3 and H4 levels were normalized to G6PDH. n=3; *\*p<0.05*, *\*\*p<0.01*, *\*\*\*p<0.001*, *\*\*\*\*p<0.0001* and *^#^p<0.05*. **(H-J)** IB analysis and quantification of histones H3 and H4 in H3WT and H3K37A *hir2Δ* normalized to G6PDH. n=4; *\*p<0.05*, *\*\*p<0.01*, *\*\*\*p<0.001*, *\*\*\*\*p<0.0001*.

To independently test this possibility, we spotted H3WT, H3K37A, and the *set2Δ* mutants to control plates, or plates containing either rapamycin, the sirtuin (class III HDAC) inhibitor nicotinamide (NAM)^48^ and class I/II HDAC inhibitor sodium butyrate (SB)^49^, or all three inhibitors combined. We found that H3K37A and H3K37A *set2Δ* showed no sensitivity to HDAC inhibition, while both were exceptionally sensitive to mTORC1 inhibition as seen above (**Fig. 4C**). Combining both mTORC1 and HDAC inhibition increased the cell death of H3K37A and H3K37A *set2Δ*, whereas H3WT and H3WT *set2Δ* were unaffected (**Fig. 4C**). Since global HDAC inhibition causes histone hyperacetylation and can increase nucleosome disassembly and/or histone exchange^50^, these data reinforce the genetic results from above suggesting that mTORC1-stress induced cell death in H3K37A is linked to histone instability.

We probed this possibility further by assessing whether disruption of known chromatin assembly pathways would cause synthetic lethality in H3K37A during mTORC1 stress that was similar to Set2 loss. In both H3WT and H3K37A, we deleted the *CAC2* gene that encodes a subunit of the chromatin assembly factor-1 (CAF-1) that deposits newly synthesized H3-H4 during replication^51^. Independently, we also deleted the *HIR2* gene, which encodes a subunit of the HIR complex that promotes replication-independent histone deposition^52^. Neither H3WT *cac2Δ* nor H3WT *hir2Δ* substantially increased rapamycin sensitivity, but disruption of either chromatin assembly pathway in H3K37A exacerbated the mTORC1-stress induced cytotoxicity similar to H3K37A *set2Δ* **(Fig. 4D**). Although disruption of either histone deposition pathway in H3WT failed to sensitize cells to mTORC1 stress, histone levels were reduced in both, while histones were decreased even further in the H3K37A background **(Fig. 4E-J).** However, this decreased histone expression alone cannot explain the increased mTORC1 stress sensitivity in H3K37A, since H3WT *set2Δ,* H3WT *cac2Δ,* and H3WT *hir2Δ* reduced histone levels but exhibited no overt mTORC1 growth defects. These findings suggest that loss of H3K37-dependent chromatin regulation, combined with defects in histone stability through disruption of histone deposition and/or transcription-linked nucleosome stability pathways, collaboratively cause cell death upon mTORC1 stress.

### Histone instability is linked to mTORC1 deregulation and sensitivity to mTORC1 stress

By analyzing ribosomal S6ph, which is a downstream readout for mTORC1 activity^15^, we previously found that mTORC1 signaling is increased in H3K37A^21^. A separate report from our group also showed that mTORC1 activity increases in *set2Δ*^22^. These studies further support a functional relationship between epigenetic signaling through the histone H3K36/K37 di-lysine motif and the mTORC1 pathway. To probe the mechanistic basis of this relationship further, we analyzed mTORC1 activity in H3WT, H3K37A, and the *set2Δ* backgrounds. As previously reported, mTORC1 activity increased in both H3K37A and H3WT *set2Δ*, while it was enhanced even further in H3K37A *set2Δ* **(Fig. 5A-B**). To confirm that the elevated mTORC1 signaling in H3K37A was due solely to H3K37A, we transformed H3WT and H3K37A with a control vector, or H3K37A with a vector expressing either *HHT2-HHF2*, *hht2K37A-HHF2*, or *hht2K37R-HHF2,* and then analyzed mTORC1 activity. While control vector H3K37A exhibited higher mTORC1 signaling compared to H3WT control, both the *HHT2-HHF2* and *hht2K37R-HHF2* vectors reduced mTORC1 activity, while expression of *hht2K37A-HHF2* had no effect **(Fig. 5C-D**). Because loss of replication-dependent (*cac2Δ*) and replication-independent (*hir2Δ*) histone deposition in H3K37A caused enhanced mTORC1 stress sensitivity and reduced histone expression, we also quantified their mTORC1 activity. Similar to H3K37A, we found that *cac2Δ* and *hir2Δ* in H3WT increased mTORC1 signaling comparable to that of H3K37A, which was not further increased in the H3K37A background **(Fig. 5E-H).** Overall, these data indicate that defects in histone stability (H3K37A), transcription-linked chromatin stability (*set2Δ*), or impaired histone deposition (*cac2Δ* or *hir2Δ*) reduce histone levels and deregulate mTORC1.

**Figure 5.**
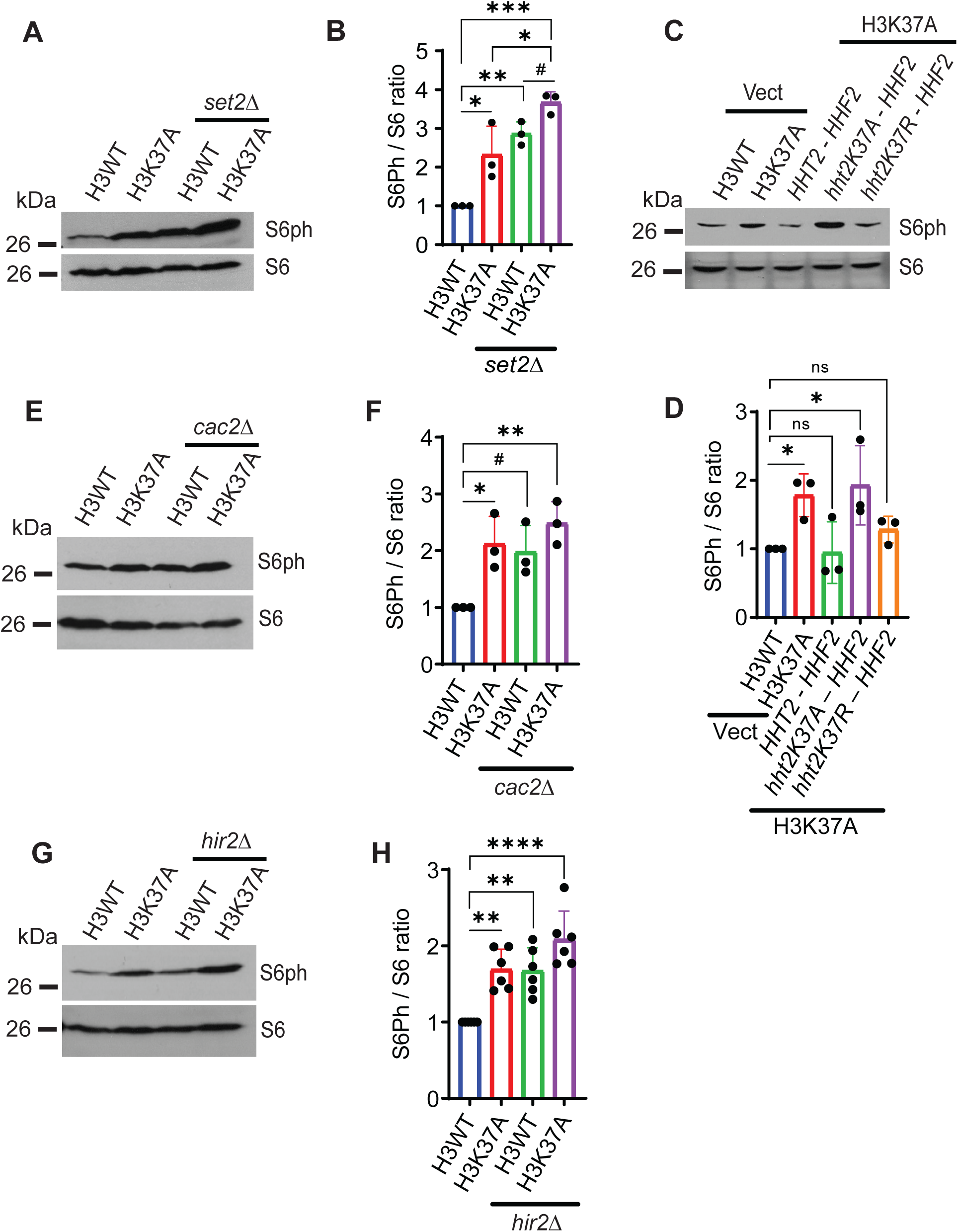
mTORC1 activity exhibits a negative correlation with the expression of histone proteins. **(A-B)** mTORC1 activity was assessed in H3WT, H3K37A, and their *set2Δ* derivatives by quantifying S6ph normalized to total S6. n=3; *\*p<0.05*, *\*\*p<0.01*, *\*\*\*p<0.001* and *^#^p<0.05*. **(C-D)** As in **A,** but mTORC1 activity was quantified in H3WT and H3K37A transformed with either control vector, or vector expressing either H3WT, H3K37A, or H3K37R. n=3; \**p<0.05*. **(E-F)** As in **A**, but mTORC1 signaling was assessed in the H3WT and H3K37A *cac2Δ* mutants. n=3; *\*p<0.05*, *\*\*p<0.01*, and *^#^p<0.05*. **(G-H)** As in **E** but mTORC1 activity was measured in the H3WT and H3K37A *cac2Δ* background. n=6; *\*\*p<0.01*, *\*\*\*\*p<0.0001*.

Tco89 is a nonessential mTORC1 subunit that recruits mTORC1 to the vacuole by interacting with the Gtr1/Gtr2 Rag GTPase subunits of the EGO complex^9^. Tco89 loss reduces mTORC1 activity by disrupting mTORC1-EGO interactions, and it results in mTORC1 repositioning from the vacuole to signaling endosomes^53,54^. To address whether increased mTORC1 activity alone was sufficient to cause cell death during mTORC1 stress, we transformed H3WT with control vector or a vector overexpressing Tco89 (*TCO89OE*) and quantified mTORC1 activity. Compared to the control vector, *TCO89OE* increased mTORC1 signaling (**Fig. 6A-B**), which is most likely explained by Tco89 enhancing mTORC1 interactions with EGO^9^. However, this increased mTORC1 signaling alone was insufficient to sensitize cells to mTORC1 stress **(Fig. 6C).** We next determined if expressing H3K37A in H3WT could deregulate mTORC1. H3WT cells were transformed with a vector expressing either *HHT2-HHF2, hht2K37A-HHF2*, or *hht2K37R-HHF2*, and the S6ph/S6 ratio was then quantified. Importantly, compared to *HHT2-HHF2* expressing cells, the *hht2K37A-HHF2* boosted mTORC1 activity, whereas *hht2K37R-HHF2* expressing cells showed no increase **(Fig. 6D-E).** These cells were then scored for growth on control media and media containing a lethal (for H3K37A) concentration of rapamycin. Expressing either *HHT2-HHF2* or *hht2K37R-HHF2* did not affect rapamycin sensitivity, while *hht2K37A-HHF2* expression caused a modest mTORC1 growth defect **(Fig. 6F).** In this experiment, the *hht2K37A-HHF2* competes with the expression of endogenous H3WT, which likely limits the overall impact H3K37A overexpression has on mTORC1 stress sensitivity. However, the results do indicate that H3K37A expression alone is sufficient to increase mTORC1 signaling and sensitize cells to mTORC1 stress. H3WT *set2Δ*, H3WT *cac2Δ*, and H3WT *hir2Δ* all have reduced histone expression and elevated mTORC1 activity, yet they do not sensitize cells to mTORC1 stress (**Fig. 5**). Therefore, while H3K37A causes these same effects, it must induce additional chromatin changes to cause cell death during mTORC1 stress.

**Figure 6.**
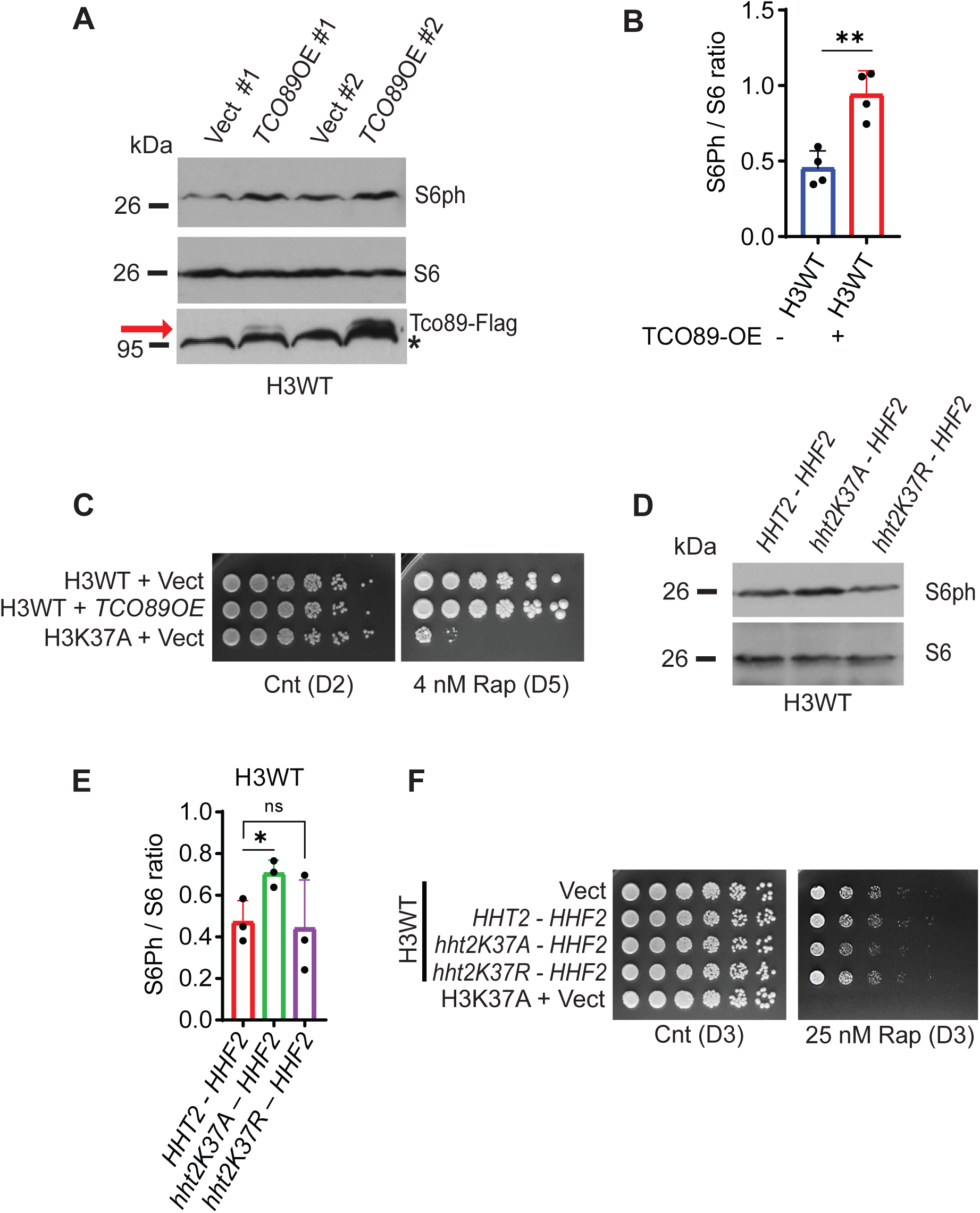
Increased mTORC1 activity alone is not responsible for cell death during mTORC1 stress. **(A-B)** H3WT was transformed with a control vector or vector expressing Tco89-FLAG (*TCO89OE*), and mTORC1 activity then was quantified by S6ph/S6 IB. Two independent transformants are shown. n=4; *\*\*p<0.01*. The red arrow indicates Tco89-FLAG, and the asterisk indicates a cross-reactive protein. **(C)** Cells from **A** were spotted onto the indicated plates for 2-5 days. **(D-E)** mTORC1 activity was measured in H3WT transformed with either *HHT2-HHF2, hht2K37A-HHF2*, or *hht2K37R-HHF2* expression vectors by quantifying S6ph/total S6. n=3; *\*p<0.05*, ns– no significance. **(F)** The cells from **D**, along with the control vector H3WT and H3K37A, were spotted onto the indicated plates for three days.

### Increased mTORC1 stress-induced cell death caused by histone instability is due to mitochondrial dysregulation

To further delineate how mTORC1 stress in H3K37A causes cell death, we isolated total RNA from H3WT and H3K37A that were either mock-treated or treated with 20 nM rapamycin for one hour and performed RNA sequencing (RNA-seq) of total RNA. By comparing each experimental condition independently to the mock-treated H3WT control, we identified differentially expressed genes (DEGs) and non-coding RNAs, including tRNAs, snRNAs, and/or snoRNAs (collectively all are referred to as DEGs), using a 1.5-fold change in expression and an FDR<0.05 as our significance criteria. **(Supplementary Table S1).** Relative to mock-treated H3WT, the mock-treated H3K37A affected 382 DEGs, the rapamycin-treated H3WT altered 140 DEGs, and the rapamycin-treated H3K37A resulted in 1070 DEGs **(Supplementary Table S1).** These DEGs were then analyzed via the STRING database to identify overrepresented cellular component gene ontologies (GOs) for each experimental condition^55^. The mock-treated H3K37A altered GOs annotated to the fungal-type cell wall, cell periphery, and anchored component of membrane categories (**Supplementary Fig. S4A**). The rapamycin-treated H3WT downregulated ribosomal gene related categories (**Supplementary Fig. S4B**), consistent with mTORC1 inhibition repressing ribosomal gene expression^56^. Interestingly, while the rapamycin-treated H3K37A altered the expression of nucleus, nucleolus, and both intracellular and cellular anatomical entity categories, ribosomal-specific GOs were not overrepresented, which is unlike the rapamycin-treated H3WT (**Supplementary Fig. S4C-D**). The individual gene lists for each condition are provided in **Supplementary Table S2.** Although the reason for this difference is not clear, one possibility could be that since H3K37A has higher basal levels of mTORC1 activity, transcriptionally repressing ribosomal genes may be delayed compared to H3WT. To determine if the increased mTORC1 signaling in H3K37A could be caused by altered expression of a known mTORC1 regulator, we manually analyzed the mock-treated H3K37A DEGs, but we did not find an established mTORC1 regulator whose expression was altered (data not shown). These data indicate that H3K37A likely does not enhance mTORC1 signaling through changes in gene expression. We also compared our DEGs to two previously published *set2Δ* transcriptomes to assess how much overlap exists between the effect that H3K37A has versus Set2 loss^22,57^. Venn analysis of these DEGs identified only 47 genes that overlap between these three studies (**Supplementary Fig. S4E** with gene lists used for comparison provided in **Supplementary Table S1**). These results further support our genetic analyses that H3K37A does not disrupt the Set2-dependent H3K36me pathway (**Fig. 3-4**).

Manual inspection of the RNA-seq dataset revealed a modest reduction (∼1.5-1.7 fold) in multiple histone genes in the mock-treated H3K37A, and also in the rapamycin-treated H3WT and rapamycin-treated H3K37A (data not shown). We independently validated this finding by RT-qPCR and found that multiple histone genes, including *HHO1,* which encodes linker histone H1, were downregulated in H3K37A compared to H3WT **(Supplementary Fig. S5A-F)**. In contrast, in **Supplementary Fig. 1F-I,** our data indicated there is no significant difference in the expression of histone genes between the control vector expressing H3WT and H3K37A cells. Currently, the explanation for these differences remains unclear, but they most likely relate to changes in the media nutrient environment required to select for plasmid maintenance in Supplementary Fig. S1F-I. We have found that nutrient-selective media required to sustain plasmid maintenance can impact mTORC1-related cell phenotypes (data not shown). Regardless, these histone mRNA decreases do not explain the reduced histone content of H3K37A, since overexpression of either *HHT2-HHF2* or *hht2K37A-HHF2* in H3K37A resulted in equivalent mRNA expression, while no difference in *HHT2* expression from the exogenous histone expression plasmid is detected **(Supplementary Fig. 1I)**. Most importantly, histone levels are restored in H3K37A specifically by proteasome inhibition **(Fig. 2D-F**), thus indicating that the histone reduction in H3K37A is due to increased histone degradation and not reduced histone mRNA repression. To specifically test if the reduction in linker histone mRNA could explain the increased mTORC1 sensitivity, we deleted *HHO1* in H3WT and H3K37A, but *HHO1* loss did not modify the mTORC1 stress response **(Supplementary Fig. S5G)**. As such, the reduction in *HHO1* mRNA does not explain the mTORC1-stress induced cell death in H3K37A.

We next performed GO analysis of the DEGs between the mock-treated H3K37A and rapamycin-treated H3K37A to identify GO categories that could explain the mTORC1-stress induced cytotoxicity. This analysis revealed that the rapamycin-treated H3K7A GOs were dominated by mitochondrial-related categories, including GOs annotated to the respiratory chain complex, oxidoreductase complex, mitochondrial respirasome, respirasome, mitochondrion, and cytochrome complex (**Fig. 7A**). Manual inspection of the genes in each of these GOs revealed that the vast majority were upregulated in the rapamycin-treated H3K37A relative to mock-treated H3K37A (data not shown). We validated this increased mitochondrial gene expression by targeted RT-qPCR analysis of some of the most overexpressed mitochondrial genes (*CRC1*, *COX26*, *FMP16*, and *FMP45*). All four genes were upregulated in rapamycin-treated H3K37A relative to the other experimental conditions **(Supplementary Fig. S6A-D)**. These data indicate that during mTORC1 stress, H3K37A specifically causes deregulation of mitochondrial gene expression, which is linked to increased cell death.

**Figure 7.**
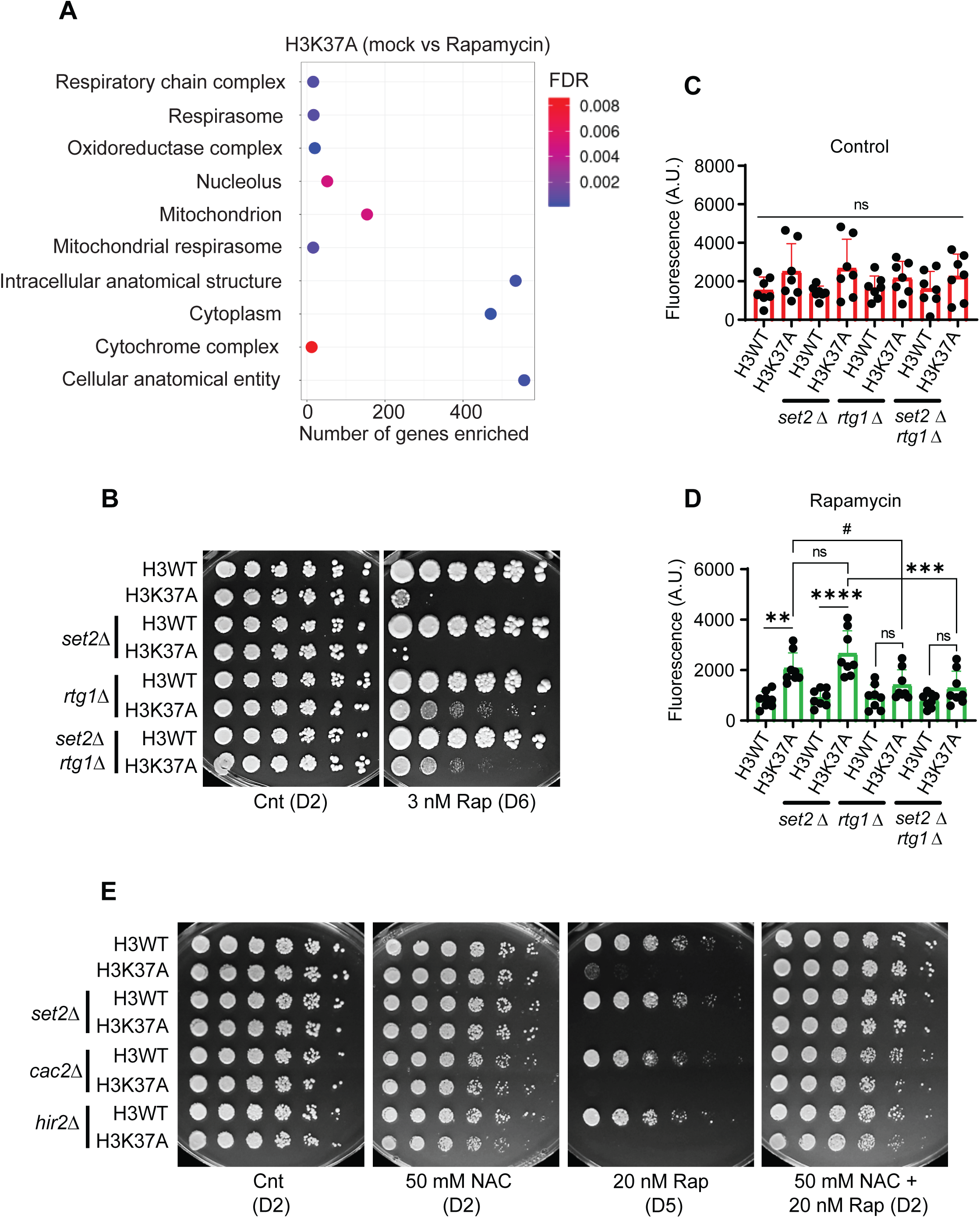
Histone instability deregulates mitochondrial retrograde signaling to cause cell death during mTORC1 stress. **(A)** Bubble blot analysis of the cellular component gene ontology (GO) categories for H3K37A (mock vs rapamycin). GO analysis was performed through the STRING database. FDR – False discovery rate. **(B)** H3WT and H3K37A combined with either *set2Δ, rtg1Δ,* or both mutations were diluted 5-fold and spotted onto the indicated plates for 2-6 days. **(C)** The indicated strains from **B** were cultured to the exponential phase and stained with DHE before quantifying DHE fluorescence. A.U.– Arbitrary Units of DHE fluorescence. n=7; ns– no significance. **(D)** As in **C**, except the cells were treated with 50 nM rapamycin for 24 hours before staining with DHE. n=8; *\*\*p<0.01*, *\*\*\*p<0.001*, *\*\*\*\*p<0.0001* and *^#^p<0.05*, ns– no significance. **(E)** H3WT and H3K37A combined with the *set2Δ, cps50Δ,* or *hir2Δ* mutations were diluted 5-fold and spotted onto control plates, or plates containing 50 mM NAC, 20 nM rapamycin, or both inhibitors for 2-5 days.

### Histone instability deregulates mitochondrial retrograde signaling to cause cell death during mTORC1 stress

The inhibition of mTORC1 leads to transcriptional activation of nuclear-encoded mitochondrial genes (retrograde signaling) through the transcription factors Rtg1 and Rtg3^58^. This retrograde signaling leads to increased mitochondrial content and enhanced mitochondrial function during nutrient stress^58^. We previously found that H3K37A increased ROS production compared to H3WT, and that the gene *CIT2*, a known retrograde-activated gene, was induced to significantly higher levels in rapamycin-treated H3K37A relative to rapamycin-treated H3WT^21^. Our RNA-seq analysis also revealed that both *CIT1* (another retrograde-activated gene) and *CIT2* genes were upregulated to higher levels in rapamycin-treated H3K37A cells compared to rapamycin-treated H3WT (**Supplementary Fig. S6E-F)**, suggesting that H3K37A may deregulate the retrograde response. To test if deregulated retrograde signaling in H3K37A causes mTORC1 stress-induced cell death, we deleted *RTG1* in H3WT, H3K37A, and their *set2Δ* derivatives to block retrograde activation. The growth of the H3WT, H3K37A, and these additional mutants was then scored on control or rapamycin-containing media. Importantly, while H3K37A and H3K37A *set2Δ* exhibited substantial cell growth defects (due to increased cell death, **Fig. 3B**), the viability of both was restored by *rtg1Δ* (**Fig. 7B**). This rescue occurred even though both H3K37A *rtg1Δ* and H3K37A *set2Δ rtg1Δ* are profoundly slow growing during mTORC1 stress compared to the matched H3WT *rtg1Δ* and H3WT *set2Δ rtg1Δ* **(Fig.7B**). These data indicate that deregulated retrograde signaling in H3K37A and H3K37A *set2Δ* causes the cytotoxic effect of mTORC1 stress.

A retrograde-mediated increase in mitochondrial activity can elevate ROS production, and we previously found evidence that mTORC1 inhibition increases ROS in H3K37A^21^. To assess if the rescue of H3K37A and H3K37A *set2Δ* viability by retrograde inactivation was due to ROS inhibition, we mock-treated or 50 nM rapamycin-treated cells for 24 hours and then quantified ROS by DHE staining. While H3K37A and H3K37A *set2Δ* increased ROS in the rapamycin-treated samples, this ROS was reduced by *rtg1Δ* (**Fig. 7C-D**). To test if deregulated ROS in mTORC1-inhibited H3K37A is the direct cause of cell death, we scored H3WT, H3K37A, and their *set2Δ* derivatives on control media, media containing either 20 nM rapamycin (which is lethal for H3K37A)^21,23^, 50 mM of the antioxidant N-acetyl cysteine (NAC), or both conditions. We also included the H3WT and H3K37A histone deposition mutants (*hir2Δ* or *cac2Δ*) to assess if mTORC1-stress induced ROS deregulation is a general phenomenon associated with histone instability. All H3WT backgrounds grew equivalently under mTORC1 stress, NAC exposure, or with both inhibitors (**Fig. 7E**). Under mTORC1 stress alone, all H3K37A backgrounds failed to grow, consistent with this level of mTORC1 stress being lethal in H3K37A; however, combined mTORC1 and ROS inhibition completely restored their viability (**Fig. 7E**). To confirm that NAC did not prevent mTORC1 inhibition by rapamycin, we assessed mTORC1 activity by analyzing S6ph/S6 and determined that rapamycin still inhibits mTORC1 in the presence of NAC (**Supplementary Fig. S6G**).

## Discussion

In this report, we demonstrate that histone H3K37 maintains chromatin stability, mTORC1 signaling regulation, and transcriptional induction of mitochondrial genes required for maintaining mitochondrial homeostasis during mTORC1 stress. Conversion of H3K37 to a small hydrophobic residue (H3K37A) increases histone instability and reduces histone levels independently of changes to histone mRNA expression or cell cycle status, indicating that the histone reduction occurs post-translationally. Indeed, proteasome inhibition fully restores histone levels in H3K37A, thus revealing the histone reduction to be due solely to increased proteasome-mediated degradation. These histone destabilizing effects were reversed by restoring an amino acid that is either positively charged (H3K37R) or polar (H3K37Q), indicating H3K37 sustains histone stability by maintaining a key histone H3 N-terminal tail contact, as we previously suggested ^21^. While H3K37 is critical for histone stability and prevents histone degradation, the reduced histone levels due to H3K37A are insufficient to increase global chromatin accessibility. This seeming incongruity is similar to the results found in a recent preprint report demonstrating that other histone destabilizing mutants also fail to increase global chromatin accessibility even though they do cause marked changes to gene-specific and lineage-specific transcriptional programs^59^. Collectively, these results indicate that cells can sustain global chromatin compaction under conditions where histone expression is reduced.

We do not yet know if H3K37A reduces Set2-mediated H3K36 methylation levels, as immunodetection methods are untrustworthy due to the underlying antibody epitope change caused by H3K37A. However, H3K37A clearly does not abolish either Set2-mediated H3K36 methylation or H3K36me reader binding since the *set2Δ*, *eaf3Δ*, and *rpd3Δ* in H3K37A cause synthetic lethality during mTORC1 stress. This possibility is further supported by the crystal structure of the Rpd3S Eaf3 chromodomain reader subunit bound to the nucleosome^34^. In this structure, H3K37 is oriented away from the H3K36me3-Eaf3 interaction where it instead contacts nucleosomal DNA, perhaps acting to stabilize H3K36me3-Eaf3 binding. Within this context, H3K37A would disrupt this stabilizing histone-DNA interaction but presumably would not interfere substantially with either Set2-mediated methylation or Eaf3-Rpd3S binding. Intriguingly, our genetic analyses of Set2 indicate that both the catalytically deficient *set2-H199L* and the SRI-truncated Set2 mutant that prevents Pol II binding partially rescue H3K37A *set2Δ* mTORC1 stress sensitivity. Both *set2-H199L* and the *set2-SRI* mutant lack endogenous H3K36me2/3, while H3K36me1 has no known epigenetic readers. As such, these results suggest that Set2 may have both methylation-dependent and methylation-independent functions in promoting a normal mTORC1 stress response. Intriguingly, a previous report found that mice deficient in the Set2 homolog SETD2 also have deregulated mTORC1 signaling^60,61^, suggesting that dysregulation of the Set2/SETD2 pathway may have evolutionarily conserved activating effects on mTORC1. Although yeast Set2 is not known to methylate non-histone substrates, SETD2 is documented to methylate some non-histone substrates^62–64^. An intriguing possibility could be that Set2, analogous to SETD2, methylates a non-histone substrate independently of Pol II binding, and this non-histone substrate contributes to the mTORC1 stress response. In support of this possibility, Set2 interacts with a diverse array of non-histone proteins as annotated in the BioGRID database, including several ribosomal proteins that would be directly relevant to mTORC1-dependent translational regulation^65^. Recent work has demonstrated that a SET domain methyltransferase can methylate ribosomal proteins^66^. An intriguing future area for investigation will be to test if Set2, and perhaps even SETD2, methylates ribosomal proteins as part of the mTORC1 pathway’s response to nutrients and/or nutrient stress.

The mTORC1 pathway is integrated into mitochondrial regulation through the retrograde signaling pathway^67^. In nutrient-rich mTORC1-activating conditions, retrograde signaling is repressed due to retention of the Rtg1/Rtg3 heterodimer in the cytoplasm. Upon mTORC1 inhibition due to nutrient insufficiency or rapamycin treatment, Rtg1/Rtg3 translocates to the nucleus and activates transcription of mitochondrial-related genes to enhance mitochondrial function^58^. We find that during mTORC1 stress the H3K37A mutant dramatically dysregulates this mitochondrial retrograde transcriptional response since mitochondrial GOs are overwhelmingly overrepresented in the rapamycin-treated H3K37A mutant compared to the H3K37A mock-treated cells. Importantly, this H3K37A mitochondrial transcriptional dysregulation ultimately is the cause of the mTORC1-stress-induced cytotoxicity since blocking retrograde activation by genetic means, or suppressing mitochondrial-derived ROS, restores H3K37A viability during mTORC1 stress. This H3K37A mitochondrial dysregulation is exacerbated further by additional chromatin destabilizing mutants, including loss of Set2 or the histone deposition complexes CAF-1 or HIR. The exact details explaining this transcriptional deregulation have yet to be established. However, one possible explanation may involve nucleosome stability at the promoters of these mitochondrial genes. Constitutively expressed (housekeeping) genes have a canonical nucleosome-free, or nucleosome-depleted, promoter structure. However, many stress-induced genes, including metabolic stress genes, have nucleosomes positioned in their promoters that must be remodeled for full transcriptional induction^68^. A previous report found that mutants that disrupt chromatin structure also deregulate mitochondrial genes in nutrient-rich conditions where these genes would normally be repressed^69^. Since our data indicate that H3K37A, as well as H3K37A Set2, CAF-1, and HIR mutants, cause chromatin instability, these promoter nucleosomes at mitochondrial genes may either be reduced or fragile and more easily remodeled upon mTORC1 stress. Such chromatin-dependent promoter changes could cause deregulation of the mitochondrial transcriptional response after mTORC1 inhibition. Because retrograde activation in wild-type cells does not normally induce cytotoxicity, this retrograde transcriptional induction in the chromatin destabilizing mutants may be uneven and result in imbalanced mitochondrial gene expression. Such altered stoichiometries of mitochondrial components ultimately could cause mitochondrial metabolic stress and ROS production. Indeed, unbalanced expression of oxidative phosphorylation pathway subunits previously was shown to increase mitochondrial ROS and reduce chronological longevity^70^. Future analysis of this mitochondrial transcriptional control in these chromatin destabilizing mutants will be required to understand how they result in mitochondrial transcriptional deregulation.

The observed increase in mTORC1 activity in H3K37A appears to be a consequence of general chromatin destabilization, since we find this occurs not just in the H3K37A and *set2Δ* as we previously reported^21,22^, but also in the CAF-1 and HIR mutants. Inspection of the mock-treated H3K37A transcriptome data did not reveal gene expression changes that readily explain the elevated mTORC1 activity. Instead, one possible explanation for this enhanced mTORC1 activity could be that increased histone instability, and the resulting histone degradation by the proteasome, boosts intracellular free amino acid levels. These free amino acids could then either be funneled into the vacuole to signal for increased mTORC1 activation by the EGO and V-ATPase complexes, or they could activate mTORC1 through less well-characterized, vacuole-independent pathways^71^. Because of the large complement of histones needed for genome packaging, it is not unreasonable to speculate that even a small percentage increase in chromatin instability-mediated histone degradation boosts intracellular amino acid levels sufficiently to enhance mTORC1 activity. Although mTORC1 is a known epigenetic regulator, such a mechanism would imply that changes to the epigenome that cause histone destabilization also could signal back to cause a corresponding increase in mTORC1 activity. Future efforts will need to pursue this possibility further to confirm these predictions.

Chromatin destabilizing histone mutants alter gene expression or chromosome function to negatively affect cellular development and stress resilience^59^. Mitochondrial activity also appears highly sensitive to changes in chromatin stability as mutants that perturb chromatin deregulate mitochondrial function in both yeast and metazoans^69,72,73^. Many of the reported phenotypic changes that occur in histone destabilizing mutants mirror those changes that occur during aging since both worm and fly models expressing histone destabilizing mutants exhibit increased functional decline indicative of accelerated aging^59,72,73^. Yeast and human cells expressing histone destabilizing mutants also exhibit defective responses to environmental stressors^59^, further supporting the idea that chromatin stability is necessary for environmental stress responsiveness^74^. During aging, epigenome integrity deteriorates while histone expression is reduced, and these chromatin changes are known to reduce longevity^75^. Intriguingly, mTORC1 activity in many organisms increases as a function of aging, whereas pharmacological mTORC1 inhibition is a geroprotective intervention that increases longevity^76^. Although it remains to be experimentally tested, we speculate that the histone loss associated with aging-dependent epigenome deterioration may fuel this increased mTORC1 activity if the histones are degraded by the proteasome and their amino acids are then recycled into the mTORC1 activation pathway. One consequence of such aging-induced increase in mTORC1 activity would be repression of nutrient-inhibited stress responses such as autophagy. Such repression would also make aging cells less resilient to nutrient stress, which is known to reduce longevity^76^. Future experiments will determine if such mTORC1-epigenetic bidirectional communication occurs, and whether deregulation of this process contributes to aging or diseases associated with mTORC1 dysregulation, such as cancer, neurodegenerative diseases, or metabolic disorders.

## Methods

### Yeast strains, plasmids, and cell growth conditions

The YNL406 (H3WT) and YNL403 (H3K37A) strains used in this study are *ura3Δ* derivatives from the histone H3/H4 library described by Dai *et. al.*^35^, which is ultimately derived from S288C. Yeast strain engineering, including gene deletion or epitope tagging, was performed as previously described^77^. All yeast strains used in this study are listed in **Supplementary Table S3.** Unless otherwise specified, yeast strains were cultivated in either nutrient-rich YPD media (1% yeast extract, 2% peptone, and 2% glucose) or in nutrient-defined SC media formulated with 0.2% yeast nitrogen base devoid of amino acids, 0.5% ammonium sulfate, 2% dextrose, and 0.19% amino acids. Dropout media to select for plasmid maintenance was prepared by leaving out the specific nutrient. Plasmids used in this study are listed in **Supplementary Table S4**. For solid media preparation, 2% agar was incorporated into the media, and specific inhibitors, such as rapamycin, nicotinamide (NAM), sodium butyrate (SB), or N-acetyl cysteine (NAC), were added just before pouring plates. In all experiments conducted, cells were cultivated in an incubated shaker at 30°C and harvested during the mid-exponential phase (OD_600_= 0.8-1.0) as determined by using a SmartSpect^TM^ Plus spectrophotometer. All yeast growth media and reagents were obtained from Research Products International, US Biologicals, Fisher Scientific, and Millipore Sigma.

### Spot assays

Cells were cultured overnight at 30 °C in either YPD or SC media, or SC media lacking the appropriate nutrient to maintain plasmid selection. Following overnight culturing, the OD_600_ was determined, and an equivalent of OD_600_=1.0 number of cells was harvested and serially diluted at a 1:5 ratio before being spotted onto the designated plates. The plates were then incubated at 30 °C, and images were captured using a MultiDoc-It Digital Imaging System at the indicated time in days.

### Site-directed mutagenesis

We cloned the *HHT2-HHF2* genetic locus into the 2µ plasmid pRS425. The resulting *HHT2-HHF2* plasmid was then used as a template for site-directed mutagenesis as follows. Primers were designed using the online software NEBaseChangerTM to change H3K37 to H3K37A and H3K37R (see **Supplementary Table S5** for primer details). Exponential PCR using the high-fidelity NEB Q5 enzyme was performed for each primer set to create mutations at H3K37A and H3K37R. The reaction mixes included 1X Q5 reaction buffer, 0.5 µM of the forward and reverse primers, 200 µM dNTPs, 0.02 U/µL of Q5 high-fidelity DNA polymerase, 25 ng of plasmid template, and nuclease-free water up to 25 µL. The mixture was incubated at 95 °C for 5 minutes for the initial denaturation, then subjected to 25 cycles at 95 °C for 1 minute, 64 °C for 1 minute, and 72 °C for 3 minutes and 40 seconds. Final extension was performed at 72 °C for 10 minutes, and the PCR products were kept at 4 °C. To remove the template plasmid, the PCR products were treated with Kinase, Ligase, & DpnI (KLD). The KLD mix contained 1 µL of the PCR product, 1X KLD reaction buffer, 1X KLD enzyme mix, and nuclease-free water to 10 µL. Samples were mixed well by pipetting, then incubated at room temperature for five minutes. After incubation, 5 μl of the KLD mix was added to ice-thawed 50 μl of JM109 competent cells and mixed, and the mixture was then incubated on ice for 30 minutes. Samples were heat-shocked at 42 °C for 30 seconds and then placed back on ice for five minutes. Subsequently, 950 μl of room temperature LB medium was added, and the mixture was incubated at 37 °C for 60 minutes with shaking. Cells were then pelleted by centrifugation, resuspended in 100 μl of LB medium, spread onto a 100 mg/mL ampicillin plate, and then incubated overnight at 37 °C. Colonies were picked, the plasmid was purified, and then screened for positive clones by Sanger sequencing.

### RNA isolation

RNA was isolated using the hot acid phenol approach^78^. Briefly, yeast cells were grown in 10 mL of SC media to OD_600_=0.8 - 1.0, pelleted, and the supernatant discarded. Pellets were resuspended in 1 mL of ice-cold water, transferred to a clean 1.5 mL microcentrifuge tube, pelleted again, and then the supernatant was removed. Cells were resuspended in 400 μL of TES solution (10 mM Tris HCl (pH 7.5), 10 mM EDTA, and 0.5% SDS), and then 400 μL of acid phenol was added before vortexing vigorously for 10 seconds. Samples were then incubated for 45 minutes at 65 °C with occasional brief vortexing, and then they were placed on ice for five minutes before centrifuging at 4 °C at 15000×rpm for five minutes. The aqueous phase was transferred to a clean 1.5 mL microcentrifuge tube, and the acid phenol treatment was repeated. After isolating the aqueous phase from the second acid phenol extraction, 400 μL of chloroform was added, the samples were vortexed vigorously, and then centrifuged for five minutes at 15,000×rpm at 4 °C. The aqueous phase was transferred to a new tube, ethanol precipitated for 30 minutes and then pelleted at 4 °C. Pellets were washed with ice-cold 70% ethanol, pelleted again, and then resuspended in 40 μL of RNase-free water and incubated at 65 °C for 3-5 minutes to dissolve the RNA pellets. All RNAs were digested with RQ1 DNase (Promega) in a 50 μL total reaction volume, incubated at 37 °C for 45 minutes. RNA was purified by phenol/chloroform extraction, precipitated, and then resuspended in 80 µL of nuclease-free water. RNA concentrations were determined by Nanodrop analysis and then stored at −70 °C.

### RNA sequencing

RNA libraries were created using the Qiagen FastSelect RNA kit, and the libraries were quantified by Qubit and Qiaxcel RNA size selection. Libraries were then sequenced on Illumina NextSeq 2000 using 2x50 P3 chip, and 20 million reads per sample were generated. Each fastq was subjected to quality control and correction using FASTQC and FASTX trimmer. The trimmed fastq was aligned to the S288 reference genome using RNA-STAR. The SAM files were then mined for the alignment of primary counts using feature Counts. The counts are normalized using TMM normalization and Log2 transformed. The transformed data matrix was loaded into R to gather statistics and determine differential expression. The mean, variance, standard deviation, and standard error of means were calculated for each gene across each condition. A principal component analysis was performed to determine different clusters in the samples. Limma with a linear regression model was used to identify genes with log2 fold change >= 0.58 and p value < 0.05. P values were then adjusted for multiplicity using the Benjamini-Hochberg method. Only genes with an adjusted p-value < 0.05 were considered differentially expressed. These genes were used to create a heatmap with unsupervised hierarchical clustering. The gene list was loaded into String DB for pathway and functional analysis. Sequencing data have been deposited at the National Institutes of Health Gene Expression Omnibus (GEO) repository with accession number GSE312461.

### cDNA synthesis and real-time quantitative PCR (RT-qPCR)

The first-strand cDNA was synthesized using the ImProm-II™ Reverse Transcription System from Promega. Briefly, 2 µg of template RNA was reverse transcribed using a modified manufacturer’s protocol. First, the template RNA was mixed with 1 µL of 500 µg/mL oligo(dT)_15_ primer and nuclease-free water up to 5 µL and placed on ice. Samples were then preheated in a thermal cycler at 70 °C for five minutes before being transferred immediately to ice for five minutes. Subsequently, 15 µL of RT reaction mix (containing 4 µL of 5X reaction buffer, 4 µL of 25 mM MgCl2, 1 µL of 10 mM dNTP, 1 µL of 40 U/µL Ribonuclease Inhibitor, 1 µL of 160 U/µL Reverse Transcriptase, and 6 µL of nuclease-free water) was added to each sample. The total 20 µL reaction mix was incubated at 25°C for five minutes, shifted to 42 °C for 60 minutes, and then inactivate the reaction by incubating at 70°C for 15 minutes. The resulting cDNA was diluted 1:10 before use. The RT-qPCR was performed using PowerUp™ SYBR™ Green Master Mix according to the manufacturer’s protocol. Briefly, the total reaction volume of 10 µL was prepared with 1 µL of cDNA, 0.75 µL of 5 µM forward and reverse primer mix, 5 µL of SYBR Green, and 4.25 µL of nuclease-free water. Amplification was performed using the StepOnePlus PCR system (Applied Biosystems). The thermal cycling conditions involved UDG activation at 50 °C for two minutes, followed by activation of dual-lock DNA polymerase at 95 °C for two minutes, and then 40 cycles of denaturation at 95 °C for three seconds and annealing/extension at 60 °C for 30 seconds. The cycle threshold (Ct) at which the fluorescence intensity crossed the threshold was used to compare samples. Target genes were normalized to the signal of the endogenous control *ACT1* (**Supplementary Table S5** for primer details). Results were expressed as fold changes in relative expression. All statistical analyses and graphs were generated using GraphPad Prism 10.

### Immunoblot (IB) analysis

The IB analysis for the proteins of interest was performed by collecting total protein from mid-log phase cells (OD_600_=0.8 - 1.0). Briefly, 50 mL of cultured cells were pelleted, washed with water, and then lysed by adding glass beads and bead beating in 120 µL of lysis buffer (10% glycerol, 10 mM Tris (pH 8.0), 0.1% NP40, 300 mM NaCl) with freshly added protease inhibitors (2 µg/mL leupeptin, 2 µg/mL pepstatin, 2 µg/mL aprotinin, 1 mM Dithiothreitol (DTT), 1x protease inhibitor cocktail and 1 mM PMSF). Lysed extracts were clarified at 4°C for 15 minutes, and the supernatant was then collected. The protein concentration was then quantified by Bradford assay, and protein extracts were stored at -70 °C. Samples (40 µg) were prepared in 2X sample buffer, denatured by boiling for five minutes, and then resolved by either 10% or 12% SDS-PAGE. Resolved proteins were transferred to a 0.45 µm PVDF membrane, and then the PVDF membrane was washed with 1X TBST buffer (Tris-buffered saline (TBS) and 1% Tween-20) before blocking with 5% milk/TBST for one hour at room temperature. Blots were then incubated with primary antibody to the following proteins and/or modified proteins: H3 (cat# 39763), H4 (cat# 39269), H2A (cat# 39235), H3K36me3 (cat# 61101), and H3K23Ac (cat#38131) from

Active Motif; G6PDH (cat# A9521) and FLAG-M2 (cat# F1804) from Sigma-Aldrich; RPS6/S6 (cat# ab40820) and Tubulin (cat# ab184970) from Abcam; phosphoS6 (Ser235/236) (cat# 22115) from Cell Signaling; HA (cat# SC805) from Santa Cruz; H3K36me2 (gifted from Dr. Strahl) ^44^. Primary antibodies were diluted in 5% milk/TBST buffer and incubated at 4 °C overnight with gentle shaking. The next day, the blot was washed three times with 1X TBST, incubated with HRP-conjugated secondary antibody (Mouse Anti-Rabbit peroxidase-conjugated IgG (cat# 211-032-171) and Goat Anti-Mouse Peroxidase-conjugated IgG (cat# 115-035-174) from Jackson Immuno Research) in blocking buffer at room temperature for one hour, and then the immunoblots were developed using the Immobilon™ Western Chemiluminescent HRP Substrate system. IBs were analyzed and quantified with ImageJ, and the band intensity data were used to generate graphical data using GraphPad Prism.

### Cell death analysis

The cell death analysis was performed using PI and YO as described previously with minor modifications^21^. Briefly, cells were grown in SC media overnight to an OD_600_=0.6-0.8 and then either mock-treated or treated with 50 nM rapamycin for six hours before harvesting. An OD_600_=1.0 cells were pelleted, washed with cold PBS, and then collected by centrifugation. Cell pellets were resuspended in 1 mL cold PBS, and then 5 µg/mL of PI and 5 µM of YO were added, mixed well, and incubated at room temperature for 20 minutes. Subsequently, 100,000 events of each stained cell sample were recorded on a Bio-Rad ZE5 flow cytometer using 488 nm excitation, and then data were collected from either a 530/30 bandpass nM green fluorescence emission for YO, or 610/20 bandpass nM red fluorescence emission for PI. Gating on cells was performed to exclude debris using single-color stained controls with standard compensation. The cell populations were categorized into three groups: live cells with low green fluorescence, apoptotic cells with higher green fluorescence, and dead cells with both red and green fluorescence. The cell count percentages of each population were used to generate graphical data in GraphPad Prism.

### Cell cycle analysis

Cells were grown in SC medium overnight to an OD_600_=0.8-1.0, and cell cycle analysis was performed as previously described with minor modifications^79,80^. In brief, 1.5 mL culture was transferred into a 15 mL centrifuge tube and pelleted at 3000xg for five minutes. Cell pellets were washed once with 1 mL cold MilliQ water and then pelleted again at 3000xg for five minutes. Pellets were resuspended in 1.5 mL cold MilliQ water before adding 3.5 mL of ice-cold ethanol (100%) dropwise while vortexing to prevent clumping, and then the samples were either incubated at room temperature for one hour or overnight at 4 °C. Cells were pelleted at 3000xg for five minutes, and the ethanol was discarded. Cell pellets were washed twice with 1 mL ice-cold sodium citrate buffer (50 mM sodium citrate, pH 7.2 adjusted with citric acid) by centrifuging at 3000xg for five minutes. The pellets were resuspended in 300 µL of RNase A solution (2 mg/mL of RNase A in 50 mM Tris (pH 8) and 15 mM NaCl) and then transferred to a 1.5 mL centrifuge tube before incubating at 37 °C for four hours with periodic mixing. After incubation, cells were collected by centrifuging at 3000xg for five minutes and then resuspended directly in 500 µL of freshly prepared protease solution (5 mg/mL pepsin in water with 4.5 µL/mL concentrated HCl added). The cell suspension was incubated at 37 °C for 20 minutes and then the cells were pelleted by centrifuging at 3000xg for five minutes. Cells then resuspended in 500 µL of ice-cold sodium citrate buffer and either kept on ice or at 4 °C until staining. For staining, 50 µL of a freshly prepared 1 mL SYTOX solution (1 µM SYTOX in sodium citrate buffer) was added to the cell suspension and then incubated at room temperature for 10 minutes. The tube containing the cells was transferred to ice and then sonicated for 15 seconds at low power (20% amplitude) with the tube kept on ice throughout the process. The sample was recorded as 20,000 events in the Bio-Rad ZE5 flow cytometer and analyzed using FlowJo software as previously described ^80^.

### MNase assay

The MNase assay was performed as described previously with minor modifications ^39,81^. Briefly, cells were grown in 100 mL SC media overnight and then collected at an OD_600_=0.8-1.0 by centrifuging at 3000xg for five minutes. Cell pellets were washed with 1.5 mL of cold water and then pelleted at 3000xg for five minutes before either continuing or freezing samples at -80 °C. To perform the MNase assay, cell pellets were washed in cold PSB buffer (20 mM Tris-HCl (pH 7.4), 100 mM NaCl, and 10 mM 2-β-mercaptoethanol (2-βME)), followed by cold ST buffer (1 M sorbitol and 20 mM Tris-HCl (pH 7.4)). Cell pellets were resuspended in 1.5 mL of cold ST buffer and measure the initial OD at OD_600_ by diluting 30 µL of cells in 1 mL of water, and then 10 units/mL of Zymolyase 20T in 1 mM DTT was added before incubating at 30 °C for approximately 30 minutes. Again, measure the OD_600_ by diluting 30 µL of cells in 1 mL of water. Once the OD_600_ was less than 80% of the initial starting OD_600_, a fresh ST buffer was added to the top of the tube, and spheroplasts were pelleted by centrifuging at 7500 rpm for five minutes at 4 °C. The supernatant was discarded, and the spheroplasts were then washed with 1.5 mL of cold ST buffer. Spheroplasts were pelleted again at 7500 rpm for five minutes at 4 °C, and then they were lysed in 800 µL of F-buffer, pH 6.5 (18% Ficoll-400, 40 mM K-phosphate monobasic (pH 7.0), 1 mM MgCl_2_), to which 5 mM 2-βME and protease inhibitors (10 µg/mL leupeptin, 1 µg/mL pepstatin, 1 µg/mL aprotinin, 1x protease inhibitor cocktail and 1 mM PMSF) were freshly added and then samples were incubated on ice for five minutes. Separately, a new centrifuge tube was prepared containing 600 µL FG-buffer pH 6.5 (7% Ficoll-400, 20 % glycerol, 40 mM K-phosphate monobasic (pH 7.0), 1 mM MgCl_2_, and freshly added 5 mM 2-βME and protease inhibitors. Spheroplast lysate was layered on top of the FG-buffer and then centrifuged at 12,500 rpm for 20 minutes at 4 °C. The supernatant was then discarded and the nuclear pellet then placed on ice. Nuclei were resuspended in 500 µL of MNase digestion buffer (10 mM HEPES (pH 7.5), 35 mM NaCl, 0.5 mM MgCl_2_, and 0.5 mM CaCl_2_) and divide into 5 aliquots before adding the following MNase concentrations: 0 U (undigested), 2.5 U, 5 U, 7.5 U, and 10 U. Samples were incubated at 37 °C for 10 minutes and then 12.5 µL of MNase stop solution (65 mM EDTA, 7% SDS) was added to stop the reactions. To each sample, 20 µL of 2 mg/mL RNase A was then added and incubated at 37 °C for one hour. After incubation, 20 µL of 10% SDS and ∼1 M of potassium acetate was added, and then the aqueous phase was isolated by phenol/chloroform extraction and maximum centrifugation for 10 minutes at room temperature. The purified DNA was ethanol precipitated, washed with 70% ethanol, and then resuspended in 20 µL of water. The results were visualized by resolving on a 2% agarose gel prestained with Midori green safe stain.

### Reactive oxygen species (ROS) quantification

ROS levels were quantified by DHE staining as previously described with minor modifications ^82^. Cells were grown in SC media overnight before mock treating or treating with 50 nM rapamycin for 24 hours at an OD_600_ =0.8-1.0. After treatment, an equivalent of OD_600_= 1.0 number of cells was centrifuged, washed twice with PBS, and then resuspended in 200 µL of staining buffer (0.1% glucose and 1 mM EDTA, pH 8.0, in PBS). Cells were transferred to a 96-well black flat-bottom microplate, and then 30 µM of DHE was added to each well and incubated at 30 °C for 15 minutes in the dark. ROS levels were measured using a fluorescence plate reader (BioTek Cytation 5) with an excitation wavelength of 520 nm and an emission wavelength of 600 nm ^83,84^. For experimental positive control, cells were treated with 10 mM hydrogen peroxide to increase ROS levels, while 50 mM NAC was used to confirm ROS quenching. The fluorescence readings of each group are used to generate graphical data with GraphPad Prism.

### Software and statistical analysis

All experiments were conducted with n = 3 or more independent biological replicates, as indicated in the figure legends. GraphPad Prism was used to calculate statistical significance. For comparing multiple groups, a one-way ANOVA with Bonferroni’s multiple comparisons test was performed. For comparing two groups, an unpaired t-test was used. Results are presented as the mean ± standard deviation (SD) with a 95% confidence interval, and a p-value of <0.05 was considered statistically significant. To represent the significance, we used (*) for a one-way AVONA and (^#^) for the t-test. Hypergeometric testing was used to test significance of transcriptome overlap for the data represented in **Supplementary Fig. S4E** using the online software (https://flaski.age.mpg.de/venndiagram/).

## Acknowledgements

We would like to acknowledge Dr. Liza Makowski and members of the Makowski laboratory for allowing us to use their plate reader. The graphical abstract was created using Biorender: Sigamani, V. (2026), https://BioRender.com/c2aub25.

## Funding Information

This work was supported by the National Institutes of Health [R01GM138393 to R.N.L. and R35GM126900 to B.D.S].

---

**Supplementary Figure 1.** (A) The H3WT and H3K37A cells were transformed with either a control vector or a vector expressing all four core histones, and then 5-fold serially diluted and spotted to the indicated plates for 3-8 days. **(B)** Whole cell extracts from cells in **A** were analyzed by IB for histone expression. **(C-E)** Quantification of histones H3, H4, and H2A normalized to α-tubulin. n=3-4; *\*p<0.05*, *\*\*\*\*p<0.0001*. **(F-I)** RT-qPCR analyses were performed for each of the indicated histone genes. n=3-4; ns – no significance.

**Supplementary Figure 2**. (A-B) H3WT and H3K37A were grown to mid-exponential phase, stained with SYTOX Green, and analyzed by flow cytometry. The histogram displays the SYTOX green fluorescence intensity (DNA content) on the x-axis and cell count on the y-axis. Data are representative of the cell cycle results represented in Figure 2A. **(C)** H3WT and H3K37A were treated with 2.5U, 5U, 7.5U, and 10U of MNase for 10 minutes at 37 °C before resolving samples on a 2% agarose gel. Lane 1 – 100 base pair (bp) DNA ladder, Lane 2 – 6 H3WT, Lane 7 empty, Lane 8 – 12 H3K37A. UD – Undigested.

**Supplementary Figure 3**. (A) The dual fluorescence scatter plot flow cytometry analysis of H3WT, H3K37A, and their *set2Δ* derivatives cells that were mock or 50 nM Rap-treated for 6 hours. YO PRO-1 (YO) on the X-axis and propidium iodide (PI) on the Y-axis. Data are representative for the cell death results in Figure 3B. **(B)** H3WT and H3WT *set2Δ* were transformed with the control vector, or with the vector expressing either wild-type Set2 or the indicated Set2 catalytic mutants (in H3WT *set2Δ*), and H3K36me2, H3K36me3, and total H3 were analyzed by IB. **(C)** IB analysis of histone H3K23Ac and total H3 in H3WT, H3K37A, and their *set2Δ* derivatives. **(D)** Quantification of **C**. n=3; *\*p<0.05*, *\*\*p<0.01* and *^#^p<0.05*.

**Supplementary Figure 4**. Bubble blot analysis of the cellular component gene ontology (GO) categories for the **(A)** H3WT (mock vs rapamycin), **(B)** H3WT vs H3K37A (mock), **(C)** H3WT mock vs H3K37A rapamycin, **(D)** H3WT rapamycin vs H3K37A rapamycin. GO analysis was performed through the STRING database. FDR – False discovery rate. **(E)** Venn analysis of differentially expressed genes (DEGs) from H3WT vs H3K37A (current study), WT vs *set2Δ* (McDaniel *et al.,* 2017), and WT vs *set2Δ* (Li *et al.,* 2023). Hypergeometric testing was used to analyze significance using a population size of 6275 total genes.

**Supplementary Figure 5**. (A-F) RT-qPCR analysis for the indicated histone genes. n=6; *\*p<0.05*, *\*\*p<0.01*, *\*\*\*p<0.001*, *\*\*\*\*p<0.0001*, and *^#^p<0.05*, *^##^p<0.01*, ns – no significance. **(G)** The H3WT, H3K37A, and the *hho1Δ* derivatives of each were serially diluted 5-fold and spotted to the indicated plates for four days.

**Supplementary Figure 6**. (A-D) RT-qPCR analysis of the indicated overexpressed mitochondrial genes from the H3K37A+rapamycin sample as identified by RNA-seq. n=3; *\*p<0.05*, *\*\*p<0.01*, *\*\*\*p<0.001*, ns – no significance. **(E-F)** The graphical data of log2-normalized counts for *CIT1* and *CIT2* from the RNA-seq analysis. n=3 – 4; *\*p<0.05*, *\*\*p<0.01*, *\*\*\*p<0.001*, *\*\*\*\*p<0.0001*, ns – no significance. **(G)** H3WT was either mock-treated or treated with 20 nM rapamycin, 50 mM NAC, or both inhibitors for one hour before analyzing mTORC1 activity by S6ph/S6 IB.

## Notes

### Competing Interest Statement

The authors have declared no competing interest.

## References

1. Liu, G.Y., and Sabatini, D.M. (2020). mTOR at the nexus of nutrition, growth, ageing and disease. Nat Rev Mol Cell Biol 21, 183–203. 10.1038/s41580-019-0199-y.

2. Benjamin, D., and Hall, M.N. (2017). mTORC1 Controls Synthesis of Its Activator GTP. Cell Rep 19, 2643–2644. 10.1016/j.celrep.2017.06.032.

3. Dibble, C.C., and Manning, B.D. (2013). Signal integration by mTORC1 coordinates nutrient input with biosynthetic output. Nature cell biology 15, 555–564.

4. Dazert, E., and Hall, M.N. (2011). mTOR signaling in disease. Current opinion in cell biology 23, 744–755.

5. Laplante, M., and Sabatini, D.M. (2012). mTOR signaling in growth control and disease. cell 149, 274–293.

6. Loewith, R., Jacinto, E., Wullschleger, S., Lorberg, A., Crespo, J.L., Bonenfant, D., Oppliger, W., Jenoe, P., and Hall, M.N. (2002). Two TOR complexes, only one of which is rapamycin sensitive, have distinct roles in cell growth control. Mol Cell 10, 457–468. 10.1016/s1097-2765(02)00636-6.

7. Reinke, A., Anderson, S., McCaffery, J.M., Yates, J., 3rd, Aronova, S., Chu, S., Fairclough, S., Iverson, C., Wedaman, K.P., and Powers, T. (2004). TOR complex 1 includes a novel component, Tco89p (YPL180w), and cooperates with Ssd1p to maintain cellular integrity in Saccharomyces cerevisiae. J Biol Chem *279*, 14752-14762. 10.1074/jbc.M313062200.

8. Li, S.C., and Kane, P.M. (2009). The yeast lysosome-like vacuole: endpoint and crossroads. Biochim Biophys Acta 1793, 650–663. 10.1016/j.bbamcr.2008.08.003.

9. 9. Binda, M., Peli-Gulli, M.P., Bonfils, G., Panchaud, N., Urban, J., Sturgill, T.W., Loewith, R., and De Virgilio, C. (2009). The Vam6 GEF controls TORC1 by activating the EGO complex. Mol Cell 35, 563–573. 10.1016/j.molcel.2009.06.033.

10. Nicastro, R., Peli-Gulli, M.P., Caligaris, M., Jaquenoud, M., Dokladal, L., Alba, J., Tripodi, F., Pillet, B., Brunner, M., Stumpe, M., et al. (2025). TORC1 autonomously controls its spatial partitioning via the Rag GTPase tether Tco89. Cell Rep 44, 115683. 10.1016/j.celrep.2025.115683.

11. Dechant, R., Saad, S., Ibanez, A.J., and Peter, M. (2014). Cytosolic pH regulates cell growth through distinct GTPases, Arf1 and Gtr1, to promote Ras/PKA and TORC1 activity. Mol Cell *55*, 409-421. 10.1016/j.molcel.2014.06.002.

12. Urban, J., Soulard, A., Huber, A., Lippman, S., Mukhopadhyay, D., Deloche, O., Wanke, V., Anrather, D., Ammerer, G., Riezman, H., et al. (2007). Sch9 is a major target of TORC1 in Saccharomyces cerevisiae. Mol Cell 26, 663–674. 10.1016/j.molcel.2007.04.020.

13. Huber, A., Bodenmiller, B., Uotila, A., Stahl, M., Wanka, S., Gerrits, B., Aebersold, R., and Loewith, R. (2009). Characterization of the rapamycin-sensitive phosphoproteome reveals that Sch9 is a central coordinator of protein synthesis. Genes Dev 23, 1929–1943. 10.1101/gad.532109.

14. Kamada, Y., Funakoshi, T., Shintani, T., Nagano, K., Ohsumi, M., and Ohsumi, Y. (2000). Tor-mediated induction of autophagy via an Apg1 protein kinase complex. J Cell Biol 150, 1507–1513. 10.1083/jcb.150.6.1507.

15. González, A., Shimobayashi, M., Eisenberg, T., Merle, D.A., Pendl, T., Hall, M.N., and Moustafa, T. (2015). TORC1 promotes phosphorylation of ribosomal protein S6 via the AGC kinase Ypk3 in Saccharomyces cerevisiae. PloS one 10, e0120250.

16. Laribee, R.N., and Weisman, R. (2020). Nuclear functions of TOR: impact on transcription and the epigenome. Genes 11, 641.

17. Laribee, R.N. (2018). Transcriptional and epigenetic regulation by the mechanistic target of rapamycin complex 1 pathway. Journal of molecular biology 430, 4874–4890.

18. Li, X., Egervari, G., Wang, Y., Berger, S.L., and Lu, Z. (2018). Regulation of chromatin and gene expression by metabolic enzymes and metabolites. Nat Rev Mol Cell Biol 19, 563–578. 10.1038/s41580-018-0029-7.

19. Suganuma, T., and Workman, J.L. (2018). Chromatin and Metabolism. Annu Rev Biochem 87, 27–49. 10.1146/annurev-biochem-062917-012634.

20. Dai, Z., Ramesh, V., and Locasale, J.W. (2020). The evolving metabolic landscape of chromatin biology and epigenetics. Nat Rev Genet 21, 737–753. 10.1038/s41576-020-0270-8.

21. Chen, H., Workman, J.J., Strahl, B.D., and Laribee, R.N. (2016). Histone H3 and TORC1 prevent organelle dysfunction and cell death by promoting nuclear retention of HMGB proteins. Epigenetics & chromatin 9, 34.

22. McDaniel, S.L., Hepperla, A.J., Huang, J., Dronamraju, R., Adams, A.T., Kulkarni, V.G., Davis, I.J., and Strahl, B.D. (2017). H3K36 methylation regulates nutrient stress response in Saccharomyces cerevisiae by enforcing transcriptional fidelity. Cell reports 19, 2371–2382.

23. Chen, H., Workman, J.J., Tenga, A., and Laribee, R.N. (2013). Target of rapamycin signaling regulates high mobility group protein association to chromatin, which functions to suppress necrotic cell death. Epigenetics & chromatin 6, 29.

24. Luger, K., Mader, A.W., Richmond, R.K., Sargent, D.F., and Richmond, T.J. (1997). Crystal structure of the nucleosome core particle at 2.8 A resolution. Nature 389, 251–260. 10.1038/38444.

25. Strahl, B.D., Grant, P.A., Briggs, S.D., Sun, Z.-W., Bone, J.R., Caldwell, J.A., Mollah, S., Cook, R.G., Shabanowitz, J., and Hunt, D.F. (2002). Set2 is a nucleosomal histone H3-selective methyltransferase that mediates transcriptional repression. Molecular and cellular biology 22, 1298–1306.

26. McDaniel, S.L., and Strahl, B.D. (2017). Shaping the cellular landscape with Set2/SETD2 methylation. Cellular and Molecular Life Sciences 74, 3317–3334.

27. Kizer, K.O., Phatnani, H.P., Shibata, Y., Hall, H., Greenleaf, A.L., and Strahl, B.D. (2005). A novel domain in Set2 mediates RNA polymerase II interaction and couples histone H3 K36 methylation with transcript elongation. Mol Cell Biol 25, 3305–3316. 10.1128/MCB.25.8.3305-3316.2005.

28. Carrozza, M.J., Li, B., Florens, L., Suganuma, T., Swanson, S.K., Lee, K.K., Shia, W.J., Anderson, S., Yates, J., Washburn, M.P., and Workman, J.L. (2005). Histone H3 methylation by Set2 directs deacetylation of coding regions by Rpd3S to suppress spurious intragenic transcription. Cell 123, 581–592. 10.1016/j.cell.2005.10.023.

29. Joshi, A.A., and Struhl, K. (2005). Eaf3 chromodomain interaction with methylated H3-K36 links histone deacetylation to Pol II elongation. Mol Cell 20, 971–978. 10.1016/j.molcel.2005.11.021.

30. Keogh, M.C., Kurdistani, S.K., Morris, S.A., Ahn, S.H., Podolny, V., Collins, S.R., Schuldiner, M., Chin, K., Punna, T., Thompson, N.J., et al. (2005). Cotranscriptional set2 methylation of histone H3 lysine 36 recruits a repressive Rpd3 complex. Cell 123, 593–605. 10.1016/j.cell.2005.10.025.

31. Smolle, M., Venkatesh, S., Gogol, M.M., Li, H., Zhang, Y., Florens, L., Washburn, M.P., and Workman, J.L. (2012). Chromatin remodelers Isw1 and Chd1 maintain chromatin structure during transcription by preventing histone exchange. Nat Struct Mol Biol 19, 884–892. 10.1038/nsmb.2312.

32. Gilbert, T.M., McDaniel, S.L., Byrum, S.D., Cades, J.A., Dancy, B.C., Wade, H., Tackett, A.J., Strahl, B.D., and Taverna, S.D. (2014). A PWWP domain-containing protein targets the NuA3 acetyltransferase complex via histone H3 lysine 36 trimethylation to coordinate transcriptional elongation at coding regions. Mol Cell Proteomics 13, 2883–2895. 10.1074/mcp.M114.038224.

33. Venkatesh, S., Smolle, M., Li, H., Gogol, M.M., Saint, M., Kumar, S., Natarajan, K., and Workman, J.L. (2012). Set2 methylation of histone H3 lysine 36 suppresses histone exchange on transcribed genes. Nature 489, 452–455.

34. Markert, J.W., Vos, S.M., and Farnung, L. (2023). Structure of the complete Saccharomyces cerevisiae Rpd3S-nucleosome complex. Nature Communications 14, 8128.

35. Dai, J., Hyland, E.M., Yuan, D.S., Huang, H., Bader, J.S., and Boeke, J.D. (2008). Probing nucleosome function: a highly versatile library of synthetic histone H3 and H4 mutants. Cell 134, 1066–1078.

36. Henikoff, S., and Smith, M.M. (2015). Histone variants and epigenetics. Cold Spring Harbor perspectives in biology 7, a019364.

37. Takeshige, K., Baba, M., Tsuboi, S., Noda, T., and Ohsumi, Y. (1992). Autophagy in yeast demonstrated with proteinase-deficient mutants and conditions for its induction. The Journal of cell biology 119, 301–311.

38. Johnson, D.L., Kumar, R., Kakhniashvili, D., Pfeffer, L.M., and Laribee, R.N. (2024). Ccr4-Not ubiquitin ligase signaling regulates ribosomal protein homeostasis and inhibits 40S ribosomal autophagy. Journal of Biological Chemistry 300.

39. Yu, R., Sun, L., Sun, Y., Han, X., Qin, L., and Dang, W. (2019). Cellular response to moderate chromatin architectural defects promotes longevity. Science advances 5, eaav1165.

40. Schneider, J., Wood, A., Lee, J.S., Schuster, R., Dueker, J., Maguire, C., Swanson, S.K., Florens, L., Washburn, M.P., and Shilatifard, A. (2005). Molecular regulation of histone H3 trimethylation by COMPASS and the regulation of gene expression. Mol Cell 19, 849–856. 10.1016/j.molcel.2005.07.024.

41. 41. van Leeuwen, F., Gafken, P.R., and Gottschling, D.E. (2002). Dot1p modulates silencing in yeast by methylation of the nucleosome core. Cell 109, 745–756. 10.1016/s0092-8674(02)00759-6.

42. Santos-Rosa, H., Millan-Zambrano, G., Han, N., Leonardi, T., Klimontova, M., Nasiscionyte, S., Pandolfini, L., Tzelepis, K., Bartke, T., and Kouzarides, T. (2021). Methylation of histone H3 at lysine 37 by Set1 and Set2 prevents spurious DNA replication. Molecular cell 81, 2793–2807. e2798.

43. Nguyen, A.T., and Zhang, Y. (2011). The diverse functions of Dot1 and H3K79 methylation. Genes & development 25, 1345–1358.

44. DiFiore, J.V., Ptacek, T.S., Wang, Y., Li, B., Simon, J.M., and Strahl, B.D. (2020). Unique and shared roles for histone H3K36 methylation states in transcription regulation functions. Cell Reports 31.

45. Wang, Y., Niu, Y., and Li, B. (2015). Balancing acts of SRI and an auto-inhibitory domain specify Set2 function at transcribed chromatin. Nucleic Acids Res 43, 4881–4892. 10.1093/nar/gkv393.

46. Kim, T., and Buratowski, S. (2007). Two Saccharomyces cerevisiae JmjC domain proteins demethylate histone H3 Lys36 in transcribed regions to promote elongation. J Biol Chem 282, 20827–20835. 10.1074/jbc.M703034200.

47. Guan, H., Wang, P., Zhang, P., Ruan, C., Ou, Y., Peng, B., Zheng, X., Lei, J., Li, B., and Yan, C. (2023). Diverse modes of H3K36me3-guided nucleosomal deacetylation by Rpd3S. Nature 620, 669–675.

48. Bitterman, K.J., Anderson, R.M., Cohen, H.Y., Latorre-Esteves, M., and Sinclair, D.A. (2002). Inhibition of silencing and accelerated aging by nicotinamide, a putative negative regulator of yeast sir2 and human SIRT1. J Biol Chem 277, 45099–45107. 10.1074/jbc.M205670200.

49. Steliou, K., Boosalis, M.S., Perrine, S.P., Sangerman, J., and Faller, D.V. (2012). Butyrate histone deacetylase inhibitors. BioResearch open access 1, 192–198.

50. Venkatesh, S., and Workman, J.L. (2015). Histone exchange, chromatin structure and the regulation of transcription. Nat Rev Mol Cell Biol 16, 178–189. 10.1038/nrm3941.

51. Kaufman, P.D., Kobayashi, R., and Stillman, B. (1997). Ultraviolet radiation sensitivity and reduction of telomeric silencing in Saccharomyces cerevisiae cells lacking chromatin assembly factor-I. Genes Dev 11, 345–357. 10.1101/gad.11.3.345.

52. Green, E.M., Antczak, A.J., Bailey, A.O., Franco, A.A., Wu, K.J., Yates, J.R., and Kaufman, P.D. (2005). Replication-independent histone deposition by the HIR complex and Asf1. Current biology 15, 2044–2049.

53. Reinke, A., Anderson, S., McCaffery, J.M., Yates, J., Aronova, S., Chu, S., Fairclough, S., Iverson, C., Wedaman, K.P., and Powers, T. (2004). TOR complex 1 includes a novel component, Tco89p (YPL180w), and cooperates with Ssd1p to maintain cellular integrity in Saccharomyces cerevisiae. Journal of Biological Chemistry 279, 14752–14762.

54. Nicastro, R., Péli-Gulli, M.-P., Caligaris, M., Jaquenoud, M., Dokládal, L., Alba, J., Tripodi, F., Pillet, B., Brunner, M., and Stumpe, M. (2025). TORC1 autonomously controls its spatial partitioning via the Rag GTPase tether Tco89. Cell Reports 44.

55. Szklarczyk, D., Morris, J.H., Cook, H., Kuhn, M., Wyder, S., Simonovic, M., Santos, A., Doncheva, N.T., Roth, A., and Bork, P. (2016). The STRING database in 2017: quality-controlled protein–protein association networks, made broadly accessible. Nucleic acids research, gkw937.

56. Powers, T., and Walter, P. (1999). Regulation of ribosome biogenesis by the rapamycin-sensitive TOR-signaling pathway in Saccharomyces cerevisiae. Molecular biology of the cell 10, 987–1000.

57. Li, Y.-M., Mei, Y.-C., Liu, A.-H., Wang, R.-X., Chen, R., and Du, H.-N. (2023). Gcn5-and Bre1-mediated Set2 degradation promotes chronological aging of Saccharomyces cerevisiae. Cell Reports 42.

58. Butow, R.A., and Avadhani, N.G. (2004). Mitochondrial signaling: the retrograde response. Molecular cell 14, 1–15.

59. Tanaka, H., McCauley, B.S., Guida, C., Lei, X., Li, S., Moreno, T.M., Guillotte, K.l., Chua, Z.-M., Abele, A., Lamba, A., et al. (2025). Nucleosome stability safeguards cell identity, stress resilience and healthy aging. bioRxiv, 2025.2009.2017.676776. 10.1101/2025.09.17.676776.

60. Walter, D.M., Gladstein, A.C., Doerig, K.R., Natesan, R., Baskaran, S.G., Gudiel, A.A., Adler, K.M., Acosta, J.O., Wallace, D.C., and Asangani, I.A. (2023). Setd2 inactivation sensitizes lung adenocarcinoma to inhibitors of oxidative respiration and mTORC1 signaling. Communications Biology 6, 255.

61. Li, X., Liu, C., Zhu, Y., Rao, H., Liu, M., Gui, L., Feng, W., Tang, H., Xu, J., Gao, W.Q., and Li, L. (2021). SETD2 epidermal deficiency promotes cutaneous wound healing via activation of AKT/mTOR Signalling. Cell Prolif 54, e13045. 10.1111/cpr.13045.

62. Park, I.Y., Powell, R.T., Tripathi, D.N., Dere, R., Ho, T.H., Blasius, T.L., Chiang, Y.-C., Davis, I.J., Fahey, C.C., and Hacker, K.E. (2016). Dual chromatin and cytoskeletal remodeling by SETD2. Cell 166, 950–962.

63. Chen, K., Liu, J., Liu, S., Xia, M., Zhang, X., Han, D., Jiang, Y., Wang, C., and Cao, X. (2017). Methyltransferase SETD2-mediated methylation of STAT1 is critical for interferon antiviral activity. Cell 170, 492–506. e414.

64. Yuan, H., Han, Y., Wang, X., Li, N., Liu, Q., Yin, Y., Wang, H., Pan, L., Li, L., and Song, K. (2020). SETD2 restricts prostate cancer metastasis by integrating EZH2 and AMPK signaling pathways. Cancer cell 38, 350–365. e357.

65. Oughtred, R., Rust, J., Chang, C., Breitkreutz, B.J., Stark, C., Willems, A., Boucher, L., Leung, G., Kolas, N., and Zhang, F. (2021). The BioGRID database: A comprehensive biomedical resource of curated protein, genetic, and chemical interactions. Protein Science 30, 187–200.

66. Park, J., Wu, J., Szkop, K.J., Jeong, J., Jovanovic, P., Husmann, D., Flores, N.M., Francis, J.W., Chen, Y.-J.C., and Benitez, A.M. (2024). SMYD5 methylation of rpL40 links ribosomal output to gastric cancer. Nature 632, 656–663.

67. Liu, Z., Sekito, T., Epstein, C.B., and Butow, R.A. (2001). RTG-dependent mitochondria to nucleus signaling is negatively regulated by the seven WD-repeat protein Lst8p. EMBO J 20, 7209–7219. 10.1093/emboj/20.24.7209.

68. Lai, W.K.M., and Pugh, B.F. (2017). Understanding nucleosome dynamics and their links to gene expression and DNA replication. Nat Rev Mol Cell Biol 18, 548–562. 10.1038/nrm.2017.47.

69. Galdieri, L., Zhang, T., Rogerson, D., and Vancura, A. (2016). Reduced histone expression or a defect in chromatin assembly induces respiration. Molecular and cellular biology 36, 1064–1077.

70. Bonawitz, N.D., Rodeheffer, M.S., and Shadel, G.S. (2006). Defective mitochondrial gene expression results in reactive oxygen species-mediated inhibition of respiration and reduction of yeast life span. Molecular and cellular biology 26, 4818–4829.

71. Gonzalez, A., and Hall, M.N. (2017). Nutrient sensing and TOR signaling in yeast and mammals. EMBO J 36, 397–408. 10.15252/embj.201696010.

72. Matilainen, O., Sleiman, M.S.B., Quiros, P.M., Garcia, S.M., and Auwerx, J. (2017). The chromatin remodeling factor ISW-1 integrates organismal responses against nuclear and mitochondrial stress. Nature communications 8, 1818.

73. Burkhart, K.B., Sando, S.R., Corrionero, A., and Horvitz, H.R. (2020). H3. 3 nucleosome assembly mutants display a late-onset maternal effect. Current Biology 30, 2343–2352. e2343.

74. Peters, A., Nawrot, T.S., and Baccarelli, A.A. (2021). Hallmarks of environmental insults. Cell 184, 1455–1468. 10.1016/j.cell.2021.01.043.

75. Pal, S., and Tyler, J.K. (2016). Epigenetics and aging. Sci Adv 2, e1600584. 10.1126/sciadv.1600584.

76. Mannick, J.B., and Lamming, D.W. (2023). Targeting the biology of aging with mTOR inhibitors. Nat Aging 3, 642–660. 10.1038/s43587-023-00416-y.

77. Janke, C., Magiera, M.M., Rathfelder, N., Taxis, C., Reber, S., Maekawa, H., Moreno-Borchart, A., Doenges, G., Schwob, E., and Schiebel, E. (2004). A versatile toolbox for PCR-based tagging of yeast genes: new fluorescent proteins, more markers and promoter substitution cassettes. Yeast 21, 947–962.

78. Collart, M.A., and Oliviero, S. (2001). Preparation of yeast RNA. Curr Protoc Mol Biol Chapter 13, Unit13 12. 10.1002/0471142727.mb1312s23.

79. Haase, S.B., and Reed, S.I. (2002). Improved flow cytometric analysis of the budding yeast cell cycle. Cell cycle 1, 117–121.

80. Rosebrock, A.P. (2017). Analysis of the budding yeast cell cycle by flow cytometry. Cold Spring Harbor Protocols 2017, pdb. prot088740.

81. Cole, H.A., Howard, B.H., and Clark, D.J. (2012). Genome-wide mapping of nucleosomes in yeast using paired-end sequencing. In Methods in enzymology, (Elsevier), pp. 145–168.

82. Sadowska-Bartosz, I., Pączka, A., Mołoń, M., and Bartosz, G. (2013). Dimethyl sulfoxide induces oxidative stress in the yeast Saccharomyces cerevisiae. FEMS yeast research 13, 820–830.

83. Michalski, R., Michalowski, B., Sikora, A., Zielonka, J., and Kalyanaraman, B. (2014). On the use of fluorescence lifetime imaging and dihydroethidium to detect superoxide in intact animals and ex vivo tissues: a reassessment. Free Radical Biology and Medicine 67, 278–284.

84. Zhao, H., Kalivendi, S., Zhang, H., Joseph, J., Nithipatikom, K., Vásquez-Vivar, J., and Kalyanaraman, B. (2003). Superoxide reacts with hydroethidine but forms a fluorescent product that is distinctly different from ethidium: potential implications in intracellular fluorescence detection of superoxide. Free Radical Biology and Medicine 34, 1359–1368.

