## Supplementary Table S5 for "Chromatin stability safeguards mitochondrial homeostasis and prevents mTORC1 hyperactivation"

**Supplementary Table S5 – List of Primers**

| **Primers** | **Sequencing ( 5`- 3`)** |
| --- | --- |
| qHHTS for | GCTTTGAGAGAAATCAGAAGATTCC |
| qHHTS Rev | GCAGCCAAGTTGGTGTCTTCGA |
| CYC13'UTRpcr | AATAAATAGGGACCTAGACTTCAGG |
| ADH13'UTRpcr | AAGCAACCTGACCTACAGGAAA |
| HHF2Promseq | AAGTATATTAGGATGAGG |
| HHT2Promseq | GCTATGGCTCGGTGTCAA |
| qFMP16 for | GCCGCCTTTCCTCACCTA |
| qFMP16 rev | AATCTTCCATTCCAGGTTGCT |
| qFMP45 for | ACTGTGTGTTGTCCTGGTTG |
| qFMP45 rev | AGTCCAGATGAACGCAAAGT |
| qCRC1 for | ACTTTAGTGACTGCGCCAAC |
| qCRC1 rev | TGCTAAAGTGGCTAATGAACCT |
| qCOX26 for | TTCAGCCAAGTTTTGAGATCTTC |
| qCOX26 rev | ACGCTCAATACTGCACTCCA |
| qHHO1 for | TGCCATAAAGAAGGGTGTGG |
| qHHO1 rev | GCTTGCTTGGGTTTTGGACT |
| qHTA1 for | TGTCTTGGAATATTTGGCCG |
| qHTA1 rev | TGGATGTTTGGCAAAACACC |
| qHTA2 for | GCTGTCTTAGAATATTTGGCTGC |
| qHTA2 rev | GGCAACAAGTTTTGGTGAATG |
| qHTB1 for | GGTAAGAAGAGAAGCAAGGCTAGAA |
| qHTB1 rev | GACTTCTTGTTATACGCAGCCA |
| qHTB2 for | GTCGATGGTAAGAAGAGATCTAAGG |
| qHTB2 rev | GTGGATTTCTTGTTATAAGCGGC |
| qHHT2 for | CTGTTGCCTTGAGAGAAATTAGAAG |
| qHHT2 rev | GCAGCCAGATTAGTGTCTTCAAAC |
| CYC13'UTRpcr | AATAAATAGGGACCTAGACTTCAGG |
| ADH13'UTRpcr | AAGCAACCTGACCTACAGGAAA |
| HHF2Promseq | AAGTATATTAGGATGAGG |
| HHT2Promseq | GCTATGGCTCGGTGTCAA |
| H3K37A For - | TGGTGTTAAGGCGCCTCACAGATATAAGC |
| H3K37R For | TGGTGTTAAGAGGCCTCACAGATATAAGCCAG |
| H3K37A/R Rev | CCGGTAGATGGGGCG |
| SET2 5'KO | CTTTTCAAAGGGGCTTTCGA |
| SET2 3'KO | TGATAGCCCACTAATGCGGATA |
| SET2 midORF | CTTTGCATCTTTTGTACCCAAC |
| CPS50 5`KO | TCCAAACTACAGAGGGAAGATAGAG |
| CPS50 3`KO | TCACCTTCAACATTTACGTTCAGTA |
| CPS50 midORF | GTTGACCTTAGAGCATAAGTACCAG |
| DOT1 S1 | GGTCACCAGTAATTGTGCGCTTTGGTTACATTTTGTTGTACAGTAATGCGTACGCTGCAGGTCGAC |
| DOT1 S2 | CTACTTAGTTATTCATACTCATCGTTAAAAGCCGTTCAAAGTGCCTCAATCGATGAATTCGAGCTCG |
| DOT1 midORF | ATGGGATGCGTTTGAACAAC |
| DOT1 5`KO | ACGCGTCAATGAAAAGGGAC |
| DOT1 3`KO | TTTAGGTACCTGGTCCACGG |
| RPD3 5'KO | CCCGGCTCATAGTATCTATCGAAA |
| RPD3 3'KO | CATCAATTAAAGGTTTGAGAAAAGTC |
| RPD3 midORF | TGTGCCATTAAGAGACGGTATTG |
| EAF3 5`KO | AAAAGGTGGCGAAGATTGGG |
| EAF3 3`KO | TTCAGAGCTGCCATTACCCT |
| EAF3 midORF | TACTGTGCCGGTCTCAAGTT |
| RPH1 5'KO | CACTTATTTTATAGATCTGCTGGAGA |
| RPH1 3'KO | GAATGTGGGTTTTCGCGAAT |
| RPH1 midORF | ACTCGCCAAGAGGATTCTTTTAC |
| IOC4 S1 | TTGTTAACTACATTTTTCAGAACGGCGTGTCATTCTCCGATAATGCGTACGCTGCAGGTCGAC |
| IOC4 S2 | TATTGTTCAAAAGCAGAGTACATCAACTGCAATAGCAACAGGTCAATCGATGAATTCGAGCTCG |
| IOC4 midORF | AGAGTGAGGGCGGTAATAGC |
| IOC4 5`KO | TGCTGCCACTTTCTTTCATAGG |
| IOC4 3`KO | TTGGACTATCAAAGACTGCGTC |
| PDP3 S1 | TCTTTTGTGATATTAGCTATCCCTGGTTTGGTCCGCAAGTTTATGCGTACGCTGCAGGTCGAC |
| PDP3 S2 | TTGAAACGTTGATATATTGAATTGATTCAATAAGTATTATTATCAATCGATGAATTCGAGCTCG |
| PDP3 midORF | GAGACCCACAGCTAATTCGG |
| PDP3 5`KO | ACGGAAGAAACAAGGTCTCTCT |
| PDP3 3`KO | CCGGGCCTTTATCGTTTCTAAG |
| CAC2 5'KO | TCGCTGTCTAACTGATGGGCTA |
| CAC2 3'KO | AGGGTCCATTGAGCACACAAT |
| CAC2 midORF | TGTGGGAAATATACATTACTCACCC |
| HIR2 5'KO | GTGCTAAATCGTCCATACAGAGGAA |
| HIR2 3'KO | CGTTCGAGATTAAGAAGGCA |
| HIR2 midORF | TGGAATGCAAAACGAATGAC |
| RTG1 5'KO | GGAATTAACTTCAACCACAACACAG |
| RTG1 3'KO | AAGTGTTCGTATCACTACCAACGAG |
| RTG1 ORF for | TTGAGTATCATTCCCAAAGACTTC |
| HHO1 S1 | GCTTGGCAGCGAGGGAAGCAATTATAATACAACTAAAGCAACATGCGTACGCTGCAGGTCGAC |
| HHO1 S2 | TATTGCTATCACCATTGACATTCTCGTTTGGATATTCACTTTTTAATCGATGAATTCGAGCTCG |
| HHO1 midORF for | GGATCCGCAAGCAACTTTGA |
| HHO1 5`KO | TGCACAATAACAGGTTTGATGGA |
| HHO1 3`KO | ACAGTACCGCCAACAAACAC |
