## Supplementary Table S3 for "Chromatin stability safeguards mitochondrial homeostasis and prevents mTORC1 hyperactivation"

**Supplementary Table S3 – List of strains**

| **Strain** | **Genotype** | **Source** |
| --- | --- | --- |
| BY4741 | MATa *his3Δ1 leu2Δ0 met15Δ0 ura3Δ0* BY4741 | OPEN Bio |
| H3WT | MATa *his3Δ200 leu2Δ0 lys2Δ0 trp1Δ63 ura3Δ0met15Δ0 can1::MFA1pr-HIS3 hht1-hhf1::NatMX4 hht2-hhf2::[HHTS-HHFS]- ura3Δ* H3WT | OPEN Bio |
| H3K37A | MATa *his3Δ200 leu2Δ0 lys2Δ0 trp1Δ63 ura3Δ0met15Δ0 can1::MFA1pr-HIS3 hht1-hhf1::NatMX4 hht2-hhf2::[HHTS-HHFS]- ura3Δ* H3K37A | OPEN Bio |
| YNL584 | MATa *his3Δ200 leu2Δ0 lys2Δ0 trp1Δ63 ura3Δ0met15Δ0 can1::MFA1pr-HIS3 hht1-hhf1::NatMX4 hht2-hhf2::[HHTS-HHFS]- ura3Δ* H3WT *set2∆::KanMX* | This Study |
| YNL582 | MATa *his3Δ200 leu2Δ0 lys2Δ0 trp1Δ63 ura3Δ0met15Δ0 can1::MFA1pr-HIS3 hht1-hhf1::NatMX4 hht2-hhf2::[HHTS-HHFS]- ura3Δ* H3K37A *set2∆::KanMX* | This Study |
| YNL1119 | MATa *his3Δ200 leu2Δ0 lys2Δ0 trp1Δ63 ura3Δ0met15Δ0 can1::MFA1pr-HIS3 hht1-hhf1::NatMX4 hht2-hhf2::[HHTS-HHFS]- ura3Δ* H3WT *cps50∆::KanMX* | This Study |
| YNL1121 | MATa *his3Δ200 leu2Δ0 lys2Δ0 trp1Δ63 ura3Δ0met15Δ0 can1::MFA1pr-HIS3 hht1-hhf1::NatMX4 hht2-hhf2::[HHTS-HHFS]- ura3Δ* H3K37A *cps50∆::KanMX* | This Study |
| YNL1128 | MATa *his3Δ200 leu2Δ0 lys2Δ0 trp1Δ63 ura3Δ0met15Δ0 can1::MFA1pr-HIS3 hht1-hhf1::NatMX4 hht2-hhf2::[HHTS-HHFS]- ura3Δ* H3WT *dot1∆::HphNT1* | This Study |
| YNL1130 | MATa *his3Δ200 leu2Δ0 lys2Δ0 trp1Δ63 ura3Δ0met15Δ0 can1::MFA1pr-HIS3 hht1-hhf1::NatMX4 hht2-hhf2::[HHTS-HHFS]- ura3Δ* H3K37A *dot1∆::HphNT1* | This Study |
| YNL1051 | MATa *his3Δ200 leu2Δ0 lys2Δ0 trp1Δ63 ura3Δ0met15Δ0 can1::MFA1pr-HIS3 hht1-hhf1::NatMX4 hht2-hhf2::[HHTS-HHFS]- ura3Δ* H3WT *rpd3∆::KanMX* | This Study |
| YNL1053 | MATa *his3Δ200 leu2Δ0 lys2Δ0 trp1Δ63 ura3Δ0met15Δ0 can1::MFA1pr-HIS3 hht1-hhf1::NatMX4 hht2-hhf2::[HHTS-HHFS]- ura3Δ* H3K37A *rpd3∆::KanMX* | This Study |
| YNL1037 | MATa *his3Δ200 leu2Δ0 lys2Δ0 trp1Δ63 ura3Δ0met15Δ0 can1::MFA1pr-HIS3 hht1-hhf1::NatMX4 hht2-hhf2::[HHTS-HHFS]- ura3Δ* H3WT *eaf3∆::KanMX* | This Study |
| YNL1046 | MATa *his3Δ200 leu2Δ0 lys2Δ0 trp1Δ63 ura3Δ0met15Δ0 can1::MFA1pr-HIS3 hht1-hhf1::NatMX4 hht2-hhf2::[HHTS-HHFS]- ura3Δ* H3K37A *eaf3∆::KanMX* | This Study |
| YNL646 | MATa *his3Δ200 leu2Δ0 lys2Δ0 trp1Δ63 ura3Δ0met15Δ0 can1::MFA1pr-HIS3 hht1-hhf1::NatMX4 hht2-hhf2::[HHTS-HHFS]- ura3Δ* H3WT *rph1∆::KanMX* | This Study |
| YNL647 | MATa *his3Δ200 leu2Δ0 lys2Δ0 trp1Δ63 ura3Δ0met15Δ0 can1::MFA1pr-HIS3 hht1-hhf1::NatMX4 hht2-hhf2::[HHTS-HHFS]- ura3Δ* H3K37A *rph1∆::KanMX* | This Study |
| YNL1147 | MATa *his3Δ200 leu2Δ0 lys2Δ0 trp1Δ63 ura3Δ0met15Δ0 can1::MFA1pr-HIS3 hht1-hhf1::NatMX4 hht2-hhf2::[HHTS-HHFS]- ura3Δ* H3WT *ioc4∆::HphNT1* | This Study |
| YNL1149 | MATa *his3Δ200 leu2Δ0 lys2Δ0 trp1Δ63 ura3Δ0met15Δ0 can1::MFA1pr-HIS3 hht1-hhf1::NatMX4 hht2-hhf2::[HHTS-HHFS]- ura3Δ* H3K37A *ioc4∆::HphNT1* | This Study |
| YNL1151 | MATa *his3Δ200 leu2Δ0 lys2Δ0 trp1Δ63 ura3Δ0met15Δ0 can1::MFA1pr-HIS3 hht1-hhf1::NatMX4 hht2-hhf2::[HHTS-HHFS]- ura3Δ* H3WT *pdp3∆::HphNT1* | This Study |
| YNL1153 | MATa *his3Δ200 leu2Δ0 lys2Δ0 trp1Δ63 ura3Δ0met15Δ0 can1::MFA1pr-HIS3 hht1-hhf1::NatMX4 hht2-hhf2::[HHTS-HHFS]- ura3Δ* H3K37A *pdp3∆::HphNT1* | This Study |
| YNL1269 | MATa *his3Δ200 leu2Δ0 lys2Δ0 trp1Δ63 ura3Δ0met15Δ0 can1::MFA1pr-HIS3 hht1-hhf1::NatMX4 hht2-hhf2::[HHTS-HHFS]- ura3Δ* H3WT *cac2∆::KanMX* | This Study |
| YNL1271 | MATa *his3Δ200 leu2Δ0 lys2Δ0 trp1Δ63 ura3Δ0met15Δ0 can1::MFA1pr-HIS3 hht1-hhf1::NatMX4 hht2-hhf2::[HHTS-HHFS]- ura3Δ* H3K37A *cac2∆::KanMX* | This Study |
| YNL1255 | MATa *his3Δ200 leu2Δ0 lys2Δ0 trp1Δ63 ura3Δ0met15Δ0 can1::MFA1pr-HIS3 hht1-hhf1::NatMX4 hht2-hhf2::[HHTS-HHFS]- ura3Δ* H3WT *hir2∆::HphNT1* | This Study |
| YNL1257 | MATa *his3Δ200 leu2Δ0 lys2Δ0 trp1Δ63 ura3Δ0met15Δ0 can1::MFA1pr-HIS3 hht1-hhf1::NatMX4 hht2-hhf2::[HHTS-HHFS]- ura3Δ* H3K37A *hir2∆::HphNT1* | This Study |
| YNL1243 | MATa *his3Δ200 leu2Δ0 lys2Δ0 trp1Δ63 ura3Δ0met15Δ0 can1::MFA1pr-HIS3 hht1-hhf1::NatMX4 hht2-hhf2::[HHTS-HHFS]- ura3Δ* H3WT *rtg1∆::HphNT1* | This Study |
| YNL1245 | MATa *his3Δ200 leu2Δ0 lys2Δ0 trp1Δ63 ura3Δ0met15Δ0 can1::MFA1pr-HIS3 hht1-hhf1::NatMX4 hht2-hhf2::[HHTS-HHFS]- ura3Δ* H3K37A *rtg1∆::HphNT1* | This Study |
| YNL1249 | MATa *his3Δ200 leu2Δ0 lys2Δ0 trp1Δ63 ura3Δ0met15Δ0 can1::MFA1pr-HIS3 hht1-hhf1::NatMX4 hht2-hhf2::[HHTS-HHFS]- ura3Δ* H3WT *set2∆::KanMX* *rtg1∆::HphNT1* | This Study |
| YNL1251 | MATa *his3Δ200 leu2Δ0 lys2Δ0 trp1Δ63 ura3Δ0met15Δ0 can1::MFA1pr-HIS3 hht1-hhf1::NatMX4 hht2-hhf2::[HHTS-HHFS]- ura3Δ* H3K37A *set2∆::KanMX* *rtg1∆::HphNT1* | This Study |
| YNL1200 | MATa *his3Δ200 leu2Δ0 lys2Δ0 trp1Δ63 ura3Δ0met15Δ0 can1::MFA1pr-HIS3 hht1-hhf1::NatMX4 hht2-hhf2::[HHTS-HHFS]- ura3Δ* H3WT *hho1∆::HphNT1* | This Study |
| YNL1202 | MATa *his3Δ200 leu2Δ0 lys2Δ0 trp1Δ63 ura3Δ0met15Δ0 can1::MFA1pr-HIS3 hht1-hhf1::NatMX4 hht2-hhf2::[HHTS-HHFS]- ura3Δ* H3K37A *hho1∆::HphNT1* | This Study |
