## Supplementary Table S4 for "Chromatin stability safeguards mitochondrial homeostasis and prevents mTORC1 hyperactivation"

**Supplementary Table S4 – List of Plasmids**

| **Plasmid** | **Description** | **Source** |
| --- | --- | --- |
| pRS425 | *2µ; LEU2* | [1] |
| pRS426 | *2µ; URA3* | [1] |
| pCC67 | HHT1-HHF1; HTA1-HTB1 2u URA3 | [2] |
| *p*HHT2-HHF2 | Histone *HHT2-HHF2* complete gene with endogenous promoter and 3`UTR in pRS426 (using *EcoRI*, SacI site) | This Study |
| *phht2*K37A*-HHF2* | Site direct mutagenesis using pHHT2-HHF2, *hht2*K37A-*HHF2* - pRS426 backbone | This Study |
| *phht2*K37R*-HHF2* | Site direct mutagenesis using pVS1, *hht2*K37R-*HHF2* - pRS426 backbone | This Study |
| pRS415 SET2 | pRS415 WT Set2-HA | [3] |
| pJD01 | pRS415 SET2 Y149F-HA | [3] |
| pJD03 | pRS415 SET2 H199L-HA | [3] |
| pJD02 | pRS415 SET2 F234Y-HA | [3] |
| pRS415_set2(1-618)-HA | *set2* (1-618) - HA (lack of *SRI* domain of *Set2*) cloned into pRS415 | [4] |
| pADH TCO89-Flag | TCO89 open reading frame cloned as a C-terminal mono-Flag fusion into the *XbaI/BamHI* sites of pN827 | [5] |
| pN827 | *pADH1 CEN URA3* | [6] |

**Reference**

1.	Baker Brachmann, C., et al., *Designer deletion strains derived from Saccharomyces cerevisiae S288C: a useful set of strains and plasmids for PCR‐mediated gene disruption and other applications.* Yeast, 1998. **14**(2): p. 115-132.

2.	Clark-Adams, C.D., et al., *Changes in histone gene dosage alter transcription in yeast.* Genes & development, 1988. **2**(2): p. 150-159.

3.	DiFiore, J.V., et al., *Unique and shared roles for histone H3K36 methylation states in transcription regulation functions.* Cell Reports, 2020. **31**(10).

4.	Kizer, K.O., et al., *A novel domain in Set2 mediates RNA polymerase II interaction and couples histone H3 K36 methylation with transcript elongation.* Molecular and cellular biology, 2005. **25**(8): p. 3305-3316.

5.	Chen, H., et al., *The histone H3 lysine 56 acetylation pathway is regulated by target of rapamycin (TOR) signaling and functions directly in ribosomal RNA biogenesis.* Nucleic acids research, 2012. **40**(14): p. 6534-6546.

6.	Laferté, A., et al., *The transcriptional activity of RNA polymerase I is a key determinant for the level of all ribosome components.* Genes & development, 2006. **20**(15): p. 2030-2040.
